# Frog-killing chytrid fungi deploy different strategies to regulate intracellular pressure in cell types that have or lack a cell wall

**DOI:** 10.1101/2025.05.13.653819

**Authors:** Sarah Prostak, Katrina B. Velle, Lillian K. Fritz-Laylin

**Affiliations:** Department of Biology, University of Massachusetts Amherst, Amherst, MA; Department of Biology, University of Massachusetts Dartmouth, North Dartmouth, MA, 02747; Howard Hughes Medical Institute, University of Massachusetts, Amherst, MA, 01003

## Abstract

Cell morphogenesis is crucial for the physiology of animals and fungi alike. While animals typically shape their cells using the actin cytoskeleton, fungi control cell shape through polarized deposition of new cell wall material, which is inflated by intracellular osmotic “turgor” pressure. Understanding where and when these mechanisms evolved is essential for understanding the evolution of cell morphogenesis. To this end, we study chytrid fungi, which have a cell type that lacks a cell wall (the “zoospore”) and a cell type that has a cell wall (the “sporangium”). While chytrid sporangia rely on polarized cell wall growth to control shape, we previously showed that the “frog-killing” chytrid fungus *Batrachochytrium dendrobatidis* (*Bd*) uses actin to control zoospore shape. Whether either zoospores or sporangia also use intracellular pressure regulation in cell shape control remains an open question. Here, we use live-cell imaging, environmental perturbations, and small molecule inhibitors to show that *Bd* sporangia generate and maintain turgor pressure, while *Bd* zoospores use specialized organelles called contractile vacuoles to pump water out of the cell, thereby keeping internal pressure low. Because chytrid fungi diverged prior to the evolution of the Dikarya—the fungal group comprising yeast, mushrooms, and filamentous fungi—these findings suggest that turgor pressure evolved early, and that cell morphogenesis underwent a major transition during early fungal evolution. We also suggest that the last common fungal ancestor may have, like chytrid fungi, employed stage-specific strategies for cell shape control—illustrating how developmental flexibility in cellular mechanisms can serve as a wellspring of evolutionary innovation.

## INTRODUCTION

Control of cell shape is fundamental to organismal form and function, and many cells change shape during development to mediate key aspects of their physiology. Fungal pathogens, for example, actively reshape their cell walls to initiate infection and evade the host immune system.^1–3^ In turn, white blood cells and unicellular amoebae rely on dynamic shape change to hunt microbes, deforming themselves to crawl across surfaces and engulf their prey.^4–6^ Other organisms, like chytrid fungi, naturally transition between these states, with one cell type that sculpts their cell walls to facilitate infection and another that uses shape change for locomotion.^7^ The basis of this functional, developmental, and evolutionary plasticity in the mechanisms controlling cell shape remains unclear.

Fungi and their sister taxa—animals and various amoebae^8–10^—employ distinct mechanisms to control cell shape. Yeasts and filamentous fungi regulate their shape through a combination of directed deposition of cell wall material and controlled intracellular pressure, known as turgor pressure, which inflates the cell wall.^11–13^ In contrast, animal cells and amoebae lack cell walls and instead use a dynamic actin cytoskeletal network, called the “actin cortex,” which lies directly under the plasma membrane and provides a scaffold that shapes the cell from within.^14,15^ Without the external structural support of a cell wall, these cells cannot sustain high internal pressure, as it would push outward on the plasma membrane, causing the cell to burst. Thus, while fungi require high internal pressure to maintain cell shape, their sister taxa must maintain low intracellular pressure. This pressure discrepancy suggests that there must have been a major transition in intracellular pressure regulation during fungal evolution. To understand the evolution of cell shape control, we must therefore understand the evolution of intracellular pressure regulation.

How and when fungal turgor pressure evolved remains largely unexplored, as most research on turgor pressure has focused on Dikarya—the fungal group comprising yeasts, mushrooms, and filamentous fungi,^16^ all of which are consistently encased in a cell wall. In contrast, fungi that diverged prior to the dikaryotic radiation often alternate between life stages with and without a cell wall.^7,17^ For example, the “frog-killing” chytrid fungus *Batrachochytrium dendrobatidis* (*Bd*) begins life as a unicellular “zoospore” that lacks a cell wall and supports its plasma membrane using actin.^18,19^ *Bd* zoospores later develop into “sporangia” cells, which build cell walls.^19^ The ability of chytrid fungi like *Bd* to transition between animal-like and yeast-like cell types makes these species valuable models for studying the evolution of cell pressure regulation. However, how *Bd* regulates intracellular pressure in either state remains unknown.

Here, we show that *Bd* sporangia use regulated turgor pressure similar to that of Dikarya. In contrast, *Bd* zoospores maintain low intracellular pressure by pumping water out of the cell using specialized organelles called contractile vacuoles, which are commonly found in species of freshwater amoebae. This means that, while cell morphogenesis of sporangia uses cell walls and turgor pressure, zoospore morphogenesis relies on an actin cortex and low intracellular pressure. *Bd*, therefore, naturally transitions between animal-like and dikaryotic cell shape control during its development. These findings suggest that the use of turgor pressure likely evolved before the dikaryotic radiation, and that early fungi may have, like chytrid fungi, used distinct cell shape control strategies at different developmental stages. This developmental plasticity represents a key example of how cellular flexibility can drive evolutionary innovation across the tree of life.

## RESULTS

### *Bd* sporangia have regulated turgor pressure similar to that of Dikarya

To understand the evolution of cell shape control in the fungal kingdom, we must understand the development of turgor pressure. Because *Bd* sporangia are encased by cell walls, like the cells of dikaryotic fungi, we hypothesized they too maintain regulated turgor pressure. Turgor pressure is an osmotic system and its presence is typically demonstrated by cell shrinkage and recovery upon hyperosmotic shock induced by sorbitol.^16,20,21^ We therefore stained the cell walls of *Bd* sporangia using Evans Blue, then imaged cell cross-sections before and after sorbitol treatment and observed concentration-dependent cell shrinkage (**Figure 1A**). While control cells treated with media alone maintained a constant cross-sectional area (100 ± 0.61% of their original cross-sectional area, **Figure 1B**), sporangia treated with media supplemented with 300 mM sorbitol for five minutes shrank to 91 ± 0.91% of their original cross-sectional area (**Figure 1B**; p<0.0001), and sporangia in 500 mM sorbitol shrank to 79.94 ± 0.75% of their original cross-sectional area (**Figure 1B**; p<0.0001). These data represent a cross-sectional area change of approximately 10% upon 300 mM sorbitol treatment and 20% upon 500 mM sorbitol treatment. Because large-scale cell shrinkage during hyperosmotic shock has been associated with plasma membrane delamination from the cell wall in plants and other fungi,^20,22,23^ we next checked whether something similar may occur in *Bd* sporangia. We stained membranes with FM4-64 and the cell wall with calcofluor white, and again imaged sporangia cross sections after five minutes of sorbitol treatment (**Figure 1C, Video S1**). While only 2.26 ± 1.60% of control sporangia and 6.8 ± 1.31% of sporangia treated with 300mM sorbitol exhibited plasma membrane delamination (**Figure 1D**; not significant), 25.20 ± 8.45% of cells treated with 500 mM sorbitol showed clear delaminations (**Figure 1D**; p = 0.0033). Given that we are measuring a single cell cross section, these results are likely an underestimation of the impact of hyperosmotic shock on *Bd*. Taken together, these results indicate that chytrid sporangia are, like dikaryotic fungi, under turgor pressure.

**Figure 1.**
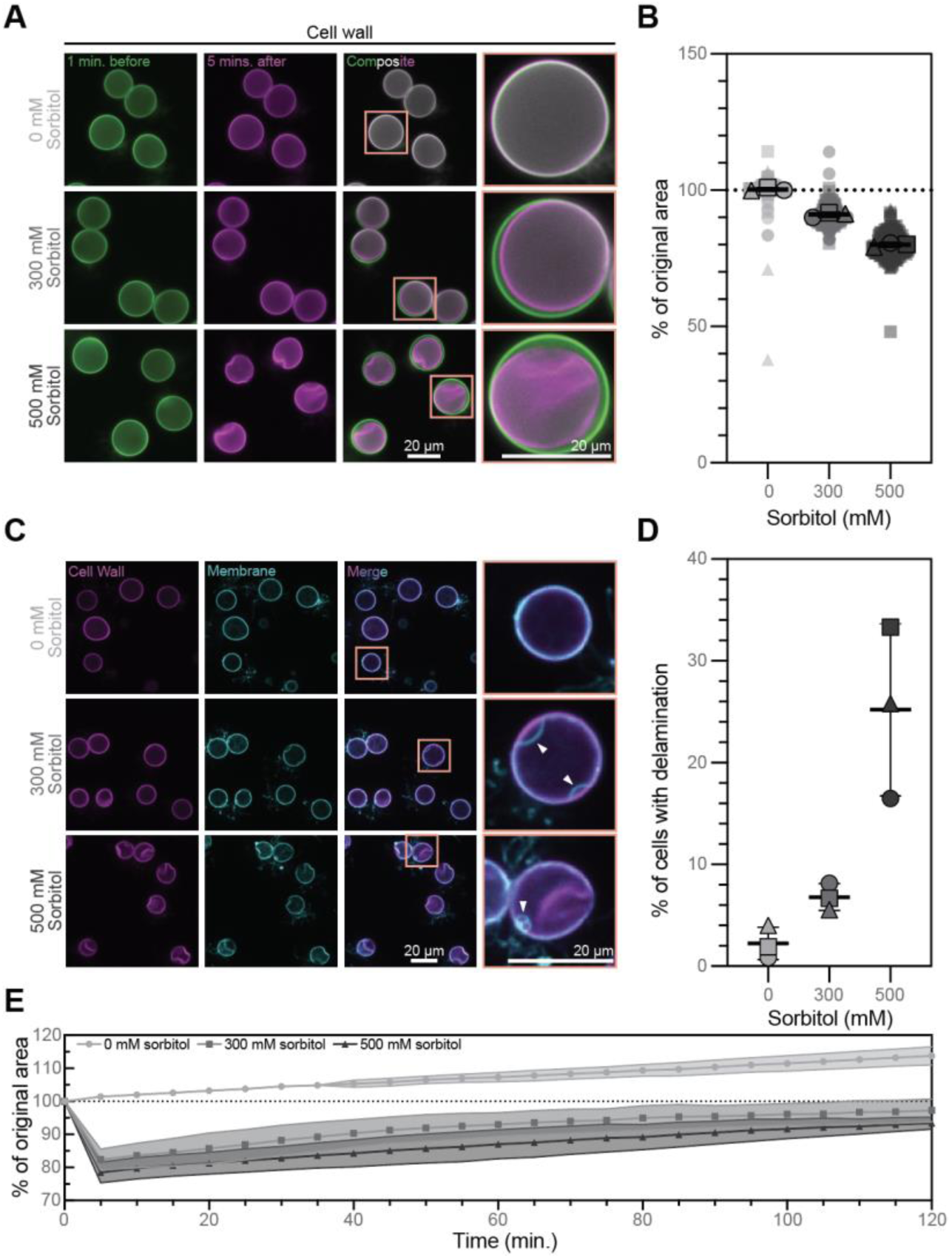
*Bd* sporangia have regulated turgor pressure. **(A)** Representative images of *Bd* sporangia with stained cell walls one minute before (green) and 5 minutes after (magenta) treatment with media supplemented with 0, 300, or 500 mM sorbitol. Composite images show the overlay of the time points. All images are adjusted to the same brightness and contrast. **(B)** Quantification of the change in cellular cross-sectional area after five minutes of the indicated sorbitol treatment, given as the percent of the cell’s cross sectional area before treatment. Each point represents one cell’s change in cross sectional area, where the average for each of three independent biological replicates is represented by a shape outlined in black. Mean and standard deviation of the replicate averages are indicated by the black lines. Statistical analysis was performed on the three replicate averages using a one-way ANOVA with Tukey’s multiple comparisons. For all comparisons: p<0.0001. **(C)** Representative images of *Bd* sporangia stained for cell walls (magenta) and membranes (cyan) after five minutes of treatment with media supplemented with 0, 300, or 500 mM sorbitol. Merged images show the overlay of membrane and cell wall. White arrowheads indicate points at which the membrane is delaminated from the cell wall. All images for each stain are adjusted to the same brightness and contrast. **(D)** Quantification of the percent of cells with clear delamination of the membrane from the cell wall in the focal plane at five minutes post treatment with the indicated sorbitol treatment. Mean and standard deviation of three independent biological replicates (black-outlined shapes) are indicated by black lines. Statistical analysis was performed on the average percentages using a one-way ANOVA with Tukey’s multiple comparisons: 0 vs. 300 n.s.; 0 vs. 500 p=0.0033; 300 vs. 500 p=0.0099. **(E)** Quantification of cellular cross-sectional area over 120 minutes of the indicated sorbitol treatment, given as the percent of the cell’s area before treatment.

Dikaryotic fungi not only possess turgor pressure, they actively regulate it in response to environmental changes.^16^ Budding yeast, for example, recover to ∼90% of their pre-stress volume within one to two hours of hyperosmotic shock.^24,25^ To determine if *Bd* also actively regulates its turgor pressure, we therefore evaluated its ability to recover volume during long-term exposure to hyperosmotic conditions. To do this, we measured the area of cross sections through *Bd* sporangia every five minutes during two hours of sorbitol treatment. While control cells steadily increased in area over the two hour imaging period, cells treated with 300 mM sorbitol shrank after five minutes to 82.3 ± 3.7% of their original area, and after two hours the cells returned to 98.3 ± 3.8% of their original area (**Figure 1E**, **Video S2**). Cells treated with 500 mM sorbitol shrank after five minutes to 78.3 ± 3.3% of their original volume, and after two hours returned to 93.7 ± 2.1% of their original volume (**Figure 1E**, **Video S2**). This represents an area increase of approximately 16% over the time of recovery for both 300 mM and 500 mM sorbitol treated cells. These findings indicate that *Bd* sporangia actively control their turgor pressure with dynamics similar to those of Dikarya.

Dikarya regulate turgor pressure primarily by controlling internal osmolyte concentrations using the high-osmolarity glycerol (HOG) signaling pathway.^16^ The HOG pathway has three main classes of proteins: 1) osmo-sensors that sense changes in osmolarity via cell wall integrity, molecular crowding, and other mechanisms;^16,21,26^ 2) a mitogen activated kinase (MAPK) cascade, ending with the MAPK Hog1, which modulates gene expression;^27–29^ and 3) signal transducers that connect the osmosensors and MAPKs. To determine whether *Bd* and other chytrids may use the HOG pathway to regulate turgor pressure, we performed a genomic survey to identify putative homologs of major HOG pathway genes in the genomes of three chytrid species–*Bd*, *Spizellomyces punctatus* (*Sp*), and *Allomyces macrogynus* (*Am*)–along with a variety of dikaryotic fungi. We found that osmosensor homolog distribution varies across chytrid species, with some having multiple putative osmosensors and others lacking any clear homolog (**Figure S2**). Furthermore, *Bd* and *Am* have the entire HOG MAPK cascade, while *Sp* appears to be missing the MAPKKKs Ssk2/22 and Ste11 (**Figure S2**). Finally, there is large variation in the repertoire of transducers among chytrid species, where *Bd* has only two of the eight investigated proteins, while *Am* has five and *Sp* has six (**Figure S2**). Taken together, these results suggest that *Bd* and other chytrids may use a version of the HOG pathway to actively regulate turgor pressure.

### *Bd* zoospores have contractile vacuoles that respond to extracellular osmolarity

Having determined that *Bd* sporangia actively maintain turgor pressure, we wondered how the zoospores that lack cell walls regulate their intracellular pressure. Because of their extreme osmotic environment, freshwater organisms that lack cell walls have evolved a variety of mechanisms to regulate internal pressure. Freshwater ciliates and amoebae, for example, use membrane-bound contractile vacuoles that literally pump water out of the cell using ion gradients to pull water into the vacuole lumen from the cytoplasm and cytoplasmic pressure to expel the water through pores connected to the extracellular environment.^30–35^ This activity can be visualized by a cycle of slow contractile vacuole growth followed by a rapid disappearance. Because contractile vacuoles have been documented in species from across the eukaryotic tree,^36–41^ we hypothesized *Bd* zoospores may use a similar mechanism to avoid bursting in their freshwater habitats. Supporting this idea, structures resembling contractile vacuoles have been observed by light and electron microscopy in the zoospores of several chytrid species,^42–45^ with the presence of these vacuoles correlated with the osmolarity of the liquid used for zoospore collection.

To explore whether chytrid zoospores possess contractile vacuoles, we searched for organelles consistent with the defining features of these structures: membrane-bound compartments whose number, size, and/or disappearance rate vary in response to external osmolarity.^36,39,42,46^ Because contractile vacuole location is highly variable, we imaged zoospores under an agarose pad to flatten the entire cell volume into more or less the same focal plane, an approach that also allows us to use contractile vacuole area as a proxy for its volume.^36^ Live-cell imaging of *Bd* zoospores under agarose using brightfield microscopy revealed small circular structures that grow and shrink largely asynchronously over time (**Figure 2A, Video S3**). To determine whether, like previously described contractile vacuoles, these structures are membrane-bound organelles, we stained *Bd* zoospores using the membrane dye FM4-64, imaged them using scanning point confocal fluorescence microscopy, and observed that these dynamic structures are indeed enclosed by membranes (**Figure 2B, Video S4**). Taken together, these results show that *Bd* has membrane-bound organelles that, like contractile vacuoles in other organisms, grow over time and then rapidly disappear.

**Figure 2.**
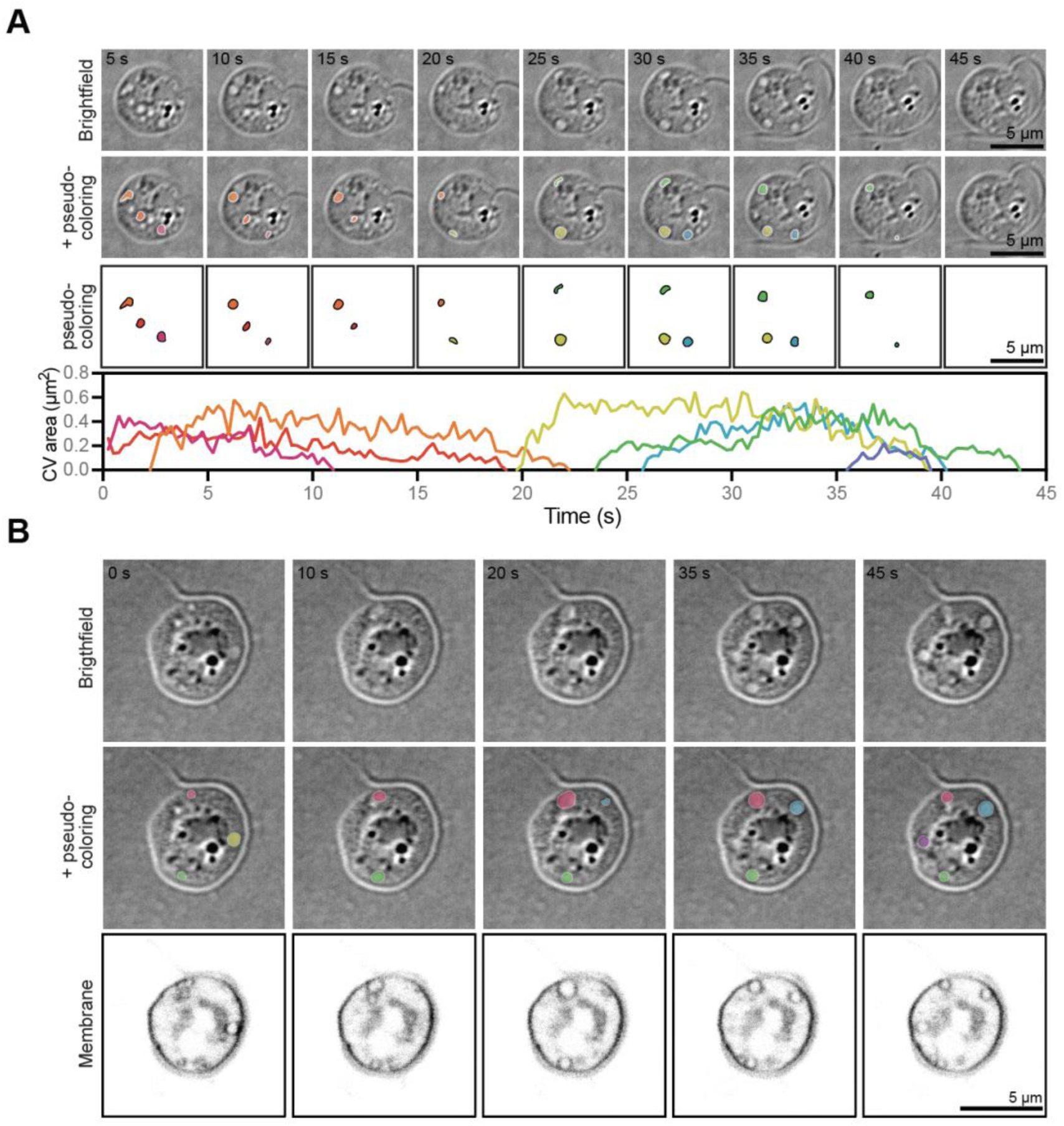
*Bd* zoospores have small, circular, membrane-bound organelles that grow and shrink over time. **(A)** Representative timelapse images of *Bd* zoospores under agarose through multiple growth and shrinkage events of small, circular organelles (top). These organelles were tracked as they grew and shrunk and are pseudo-colored by organelle (middle). The area of each organelle is graphed over time (bottom). **(B)** Representative timelapse images of membrane-stained *Bd* zoospores under agarose (bottom) with the small organelles observed in brightfield (top) pseudo-colored by organelle (middle). All membrane images are adjusted to the same brightness and contrast. All brightfield images are on an inverted LUT.

As osmoregulators, contractile vacuoles are defined by changing volume and/or pumping rate in response to changes in environmental osmolarity. In the lab, this is typically measured by observing changes in contractile vacuole activity in the presence of varying concentrations of sorbitol.^36,47^ We therefore imaged zoospores under a range of sorbitol concentrations and quantified the number, size, and dynamics of the membrane-bound organelles. We found that as sorbitol concentration increases, the average area of the largest dynamic organelle per cell decreases, starting from an average of 0.68 ± 0.22 µm^2^ in control cells treated with buffer alone, to as small as 0 ± 0.01 µm^2^ in cells treated with 100 mM sorbitol (**Figure 3A, Video S5**). The frequency of disappearance of these organelles also responded to extracellular osmolarity. The organelles in control cells averaged 6 ± 0.92 disappearance events per minute, and this rate generally decreased as sorbitol concentration increased with the organelles of cells treated with 100 mM sorbitol averaging 0.07 ± 0.12 disappearance events per minute (**Figure 3A**). Given the cell-to-cell variability of largest vacuole size and disappearance rate, both within and between biological replicates, we quantified the percent of the total cell area that is lost per minute and found again, and with slightly lower variability, that the percent of the total cell area lost generally decreased with increasing sorbitol concentrations (**Figure 3A**). These results show that the dynamic organelles observed in *Bd* respond to changes in external osmolarity, indicating that the disappearance of the vacuoles represent pumping used to regulate internal pressure, and therefore fit the definition of contractile vacuoles.

**Figure 3.**
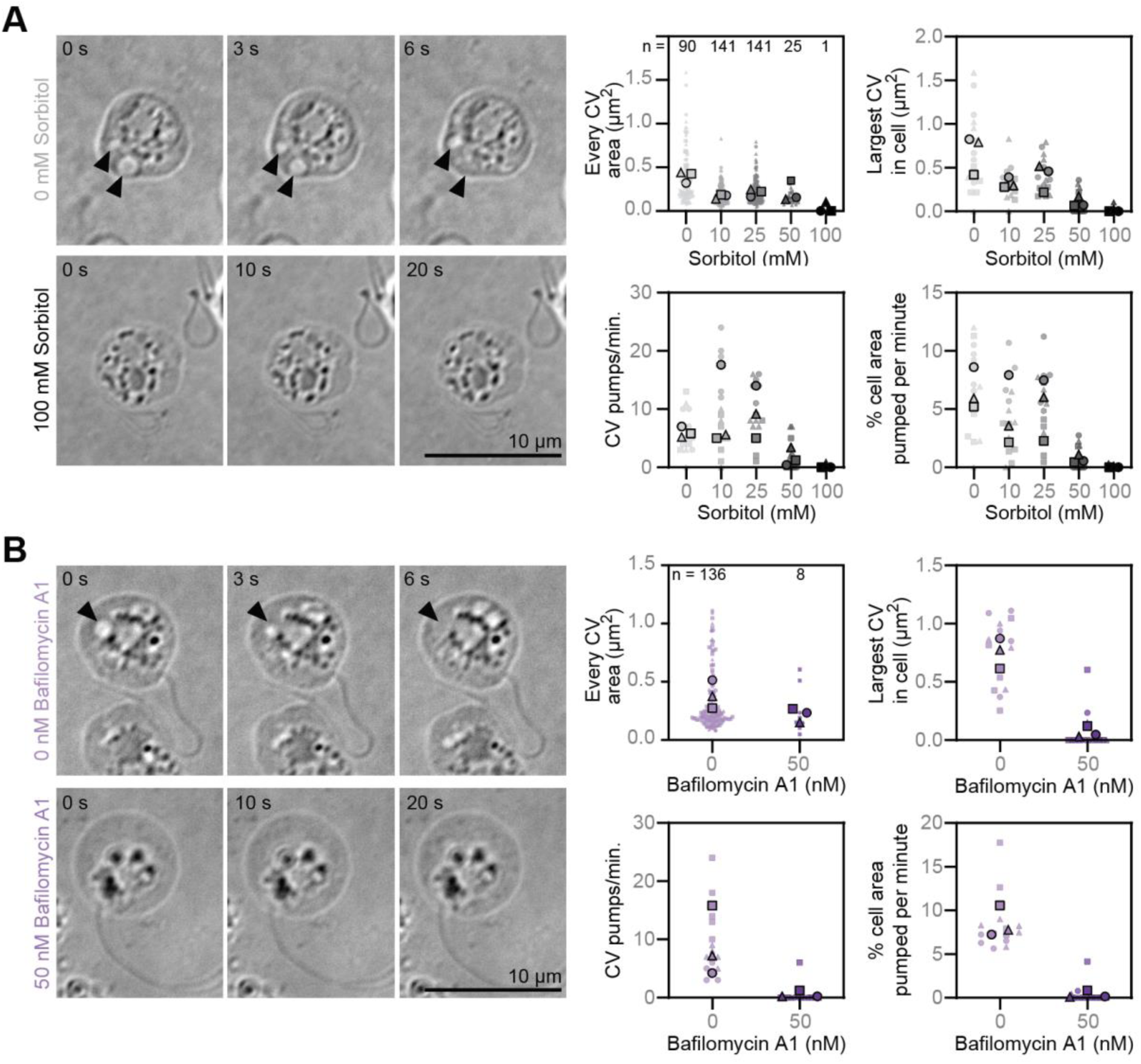
Small, membrane-bound organelles in *Bd* zoospores respond to hyperosmotic shock and rely on vacuolar-ATPase function. **(A)** Representative timelapse images of *Bd* zoospores under agarose in the given sorbitol concentrations (left) and quantification of the area, pumping rate, and percent of cell area pumped per minute of all of the putative contractile vacuoles from five random cells per sorbitol treatment (right). Black arrowheads indicate growing and shrinking organelles. Number of growing and shrinking organelles quantified indicated by n. **(B)** Representative timelapse images of *Bd* zoospores under agarose treated with or without 50 nM of the vacuolar ATPase inhibitor Bafilomycin A1 for 35 minutes (left) and quantification of vacuole area, pumping rate, and percent of cell area pumped per minute for all growing and shrinking organelles in five random cells per treatment (right). Black arrowheads indicate growing and shrinking organelles. Number of growing and shrinking organelles quantified indicated by n. All brightfield images are on an inverted LUT.

Contractile vacuole function in other species depends on vacuolar-ATPases (v-ATPases), which create an ion gradient to facilitate water flow into the contractile vacuole through aquaporins.^30–35^ To explore whether chytrids may use the same molecular mechanisms to power vacuole filling, we first searched for v-ATPases and aquaporins in the genomes of three chytrid species and found putative homologs of all subunits of the v-ATPase complex,^48,49^ along with several aquaporins (**Figure S3**). To test if v-ATPase activity is important for contractile vacuole function in *Bd*, we treated zoospores with Bafilomycin-A1, a v-ATPase inhibitor that disrupts contractile vacuole function by preventing contractile vacuole re-filling. (The predicted *Bd* v-ATPase subunits have all the conserved residues for Bafilomycin-A1 binding and function, **Figure S4A**.^50^) Treating *Bd* zoospores with 50 nM Bafilomycin-A1 for 35 minutes drastically reduced contractile vacuole size: the area of the largest contractile vacuole in Bafilomycin-A1 treated cells was 0.07 ± 0.05 µm^2^ compared to 0.75 ± 0.13 µm^2^ in cells treated with buffer alone (**Figure 3B, Video S5**). We also found that Bafilomycin-A1 treatment had drastic effects on contractile vacuole pumping, with the contractile vacuoles of Bafilomycin-A1 treated cells pumping an average of 0.53 ± 0.57 pumps per minute compared to 9.07 ± 6.02 pumps per minute in control cells (**Figure 3B**). Normalizing for cell size, we calculated that Bafilomycin-A1 treated cells pump only 0.35 ± 0.41% of their area per minute compared to control cells that pump 8.53 ± 1.79% of their area per minute (**Figure 3B**). Because v-ATPase activity is required for contractile vacuole refilling, we next quantified the number of *Bd* zoospores that undergo multiple pumping cycles in the presence of this drug. Of cells with active contractile vacuoles at the beginning of imaging, 95.5 ± 2.3% of control cells underwent multiple pump cycles compared to only 15.2 ± 11.5% of Bafilomycin-A1 treated cells (**Figure S4B**, p = 0.0007). Finally, because zoospores with nonfunctional contractile vacuoles are likely to die due to being overwatered, we quantified the percentage of cells that died over the three minute imaging period. We found that only 0.2 ± 0.21% of control cells died, while 2.42 ± 0.36% of Bafilomycin A1 treated cells died (**Figure S4C**, p= 0.0008). Taken together, these results suggest that the contractile vacuoles of *Bd* rely heavily on v-ATPase activity and are critical to survival in freshwater.

### *Bd* contractile vacuoles slow their pumping during cell wall accumulation

Having determined that *Bd* cells that lack cell walls reduce intracellular pressure, while *Bd* cells with cell walls boost it, we next wondered what happens when *Bd* transitions between these two states. Because maintaining high intracellular pressure is incompatible with a wall-less cell type, we hypothesized that this transition must involve a gradual shift from contractile vacuole activity to turgor-based pressure regulation, coordinated by regulatory mechanisms that link pressure control to cell wall assembly. To test this hypothesis, we visualized contractile vacuole activity and cell wall accumulation in cells undergoing the zoospore-to-sporangium transition—a developmental process called encystation—which can be rapidly triggered by treatment with purified mucin.^51^ We treated cells with mucin, immediately flattened them, and imaged contractile vacuole activity and cell wall accumulation after 15, 20, 25, 30, and 40 minutes (**Figure 4A**). We found that contractile vacuoles pump on average 25.53 ± 7.34 pumps per minute at 15 minutes after mucin treatment, and this rate decreases to about 1.73 ± 4.95 pumps per minute at 40 minutes after treatment (**Figure 4B**). During these 25 minutes, the total cell wall intensity increased by 1.65 ± 0.32 times (**Figure 4B**). To visualize this relationship directly, we plotted pumping rate against cell-wall intensity and found a clear trend; as cell-wall intensity increases, contractile vacuole pumping rate decreases (**Figure 4C**). We next wondered how long it takes after appreciable cell-wall accumulation for contractile vacuoles to stop pumping. To this end, we identified the first time point at which the total cell-wall intensity of an individual cell was at least 20% higher than the background, determined when that cell’s pumping rate dropped to zero, and then calculated the time difference between these two events. Although the timing varied across cells, most stopped pumping within 10–25 minutes of appreciable cell-wall accumulation (**Figure 4D**). This delay suggests that the switch from contractile vacuole activity to turgor-based pressure regulation happens gradually, likely to allow for sufficient cell-wall thickness to counter the force of turgor pressure on the membrane, and/or for the cell to produce sufficient osmolyte to modulate water influx. Either way, these two scenarios imply the existence of molecular mechanisms allowing cross talk between cell wall assembly, turgor pressure regulation, and/or contractile vacuole activity.

**Figure 4.**
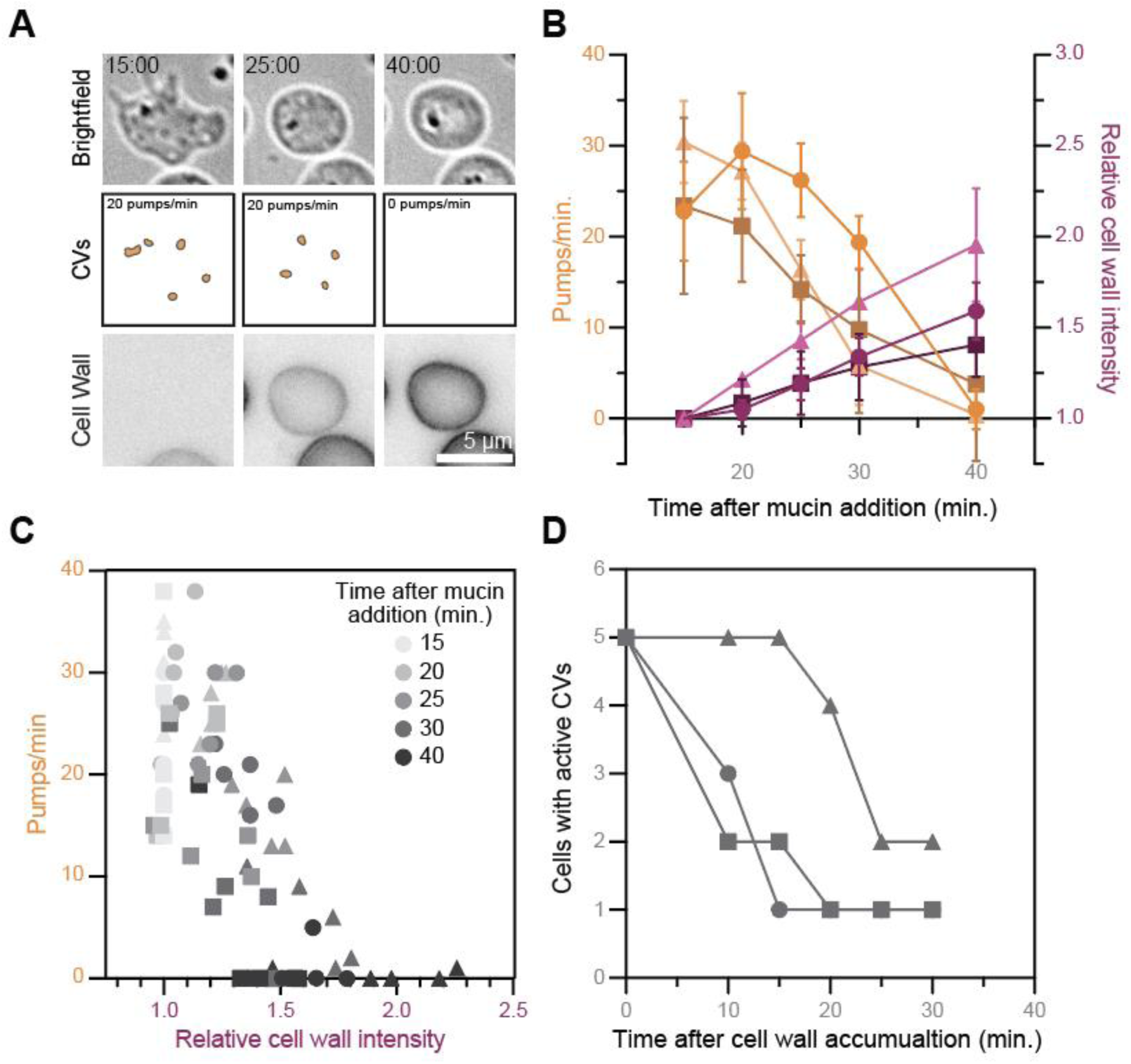
The pumping of *Bd* zoospores’ contractile vacuoles slow during cell wall assembly. **(A)** Representative time lapse images of cell-wall stained, confined *Bd* zoospores after mucin treatment to induce encystation. Contractile vacuoles are highlighted (middle). Brightfield images are on an inverted LUT. All cell wall images are adjusted to the same brightness and contrast. **(B)** Quantification of the contractile vacuole pumping rate (shades of orange, left axis) and the relative cell wall intensity (shades of magenta, right axis) over time after mucin treatment. Mean and standard deviation values for each time point for three independent replicates (shapes) are shown. Some error bars are not shown, as they were smaller than the shape itself. **(C)** All cells quantified in (B), graphed by pumping rate vs relative cell wall intensity. Time after mucin treatment is indicated in shades of grey for each shape. Each shape represents data from an independent replicate. **(D)** Quantification of the number of cells with active contractile vacuoles (>0 pumps/min) over time after cell wall accumulation. Cell wall was considered to have accumulated at the first time point where total cell wall intensity was at least 20% greater than background. Each shape represents an independent replicate.

## DISCUSSION

Here, we show that different *Bd* cell types use distinct modes of intracellular pressure regulation. While sporangia that have cell walls actively maintain high turgor pressure, zoospores that lack cell walls use contractile vacuoles to maintain low intracellular pressure. Together with previous work showing that *Bd* zoospores migrate using a dynamic actin cortex,^18^ our findings suggest that chytrids naturally alternate between two fundamentally different modes of cell shape control: an animal- or amoeba-like system in the wall-less zoospore, and a canonical fungal system in the walled sporangium. This duality makes *Bd* a valuable model for studying the evolution and development of intracellular pressure regulation and cell morphogenesis.

Because chytrid fungi diverged before the dikaryotic radiation,^52^ our discovery of regulated turgor pressure in *Bd* sporangia suggests that this mechanism of pressure control arose very early in fungal evolution and may have been present in the last common ancestor of fungi. The importance of turgor pressure for morphogenesis in Dikarya^11,12,53^ makes chytrids an important point of comparison for understanding the evolution of multicellular fungi. Importantly, the ability of *Bd* to toggle between shape control strategies associated with either animals or fungi suggests that the last common ancestor of fungi may have deployed similar context-dependent mechanisms. This finding implies that, rather than a linear evolutionary shift from amoeboid to turgor-based morphogenesis, the evolution of fungal cell shape control may have involved the co-option and reconfiguration of pre-existing systems.

In addition to shedding light on the evolution of cell shape control in fungi, this work advances our understanding of *Bd* pathogenesis. As an amphibian pathogen, *Bd* encounters dramatically different osmotic environments over the course of its life cycle. Zoospores swim in freshwater, where the near-zero osmolarity creates a constant influx of water that must be actively managed. In contrast, sporangia develop within frog epithelial tissues, where the osmolarity is approximately 200–300 mOsM,^54,55^ resulting in a far lower osmotic gradient and reduced water influx. The ability to regulate intracellular pressure across these conditions is likely essential for *Bd’*s survival and infection success. Moreover, like other fungal pathogens,^1,56^ *Bd* sporangia may use turgor pressure during its invasion of host cells or tissues.^57^

Finally, these results may provide clues about the environmental context in which fungi evolved. Although hotly debated, the two dominant hypotheses propose that fungi originated in either marine or freshwater habitats.^58^ Our finding that *Bd* zoospores use contractile vacuoles— organelles typically associated with freshwater organisms—is consistent with the idea that early fungi evolved in aquatic environments, where regulation of intracellular pressure would have been crucial for survival. The ability of early fungi to transition between using contractile vacuoles that facilitate survival in freshwater environments and using turgor pressure/cell wall systems that enable survival in terrestrial environments may have been crucial for the conquest of land. This flexibility in cellular morphogenesis exemplifies how developmental plasticity in core cellular mechanisms can facilitate organisms’ colonization of diverse ecological niches. This principle extends beyond the fungal kingdom, as developmental flexibility in cellular mechanisms serves as a wellspring of evolutionary innovation across the tree of life.

## ACKNOWLEDGEMENTS

We would like to thank Drs. Jarrett Man and Jacob Ritz for their feedback on the manuscript. We would also like to thank the Institute for Applied Life Sciences Light Microscopy Director, Dr. James Chambers for help with troubleshooting and feedback on the manuscript. This work was primarily funded by NSF grant IOS2143464 awarded to L.K.F.-L. who is Canadian Institute for Advanced Research (CIFAR) fellow in the Fungal Kingdom: Threats and Opportunities program and an Investigator of the Howard Hughes Medical Institute. This work was also supported by NIH grant R00GM147656 awarded to K.B.V. Microscopy data acquisition and analysis was partially performed on equipment that was funded by the Massachusetts Life Science Center.

## METHODS

### Cell Culture and Cell Preparation

<u>Turgor pressure experiments</u>: *Batrachochytrium dendrobatidis* JEL423 (*Bd*) zoospores from a culture grown at 24℃ in 1% tryptone (w/v; Sigma # T7293) were harvested by filtering through a sterile 40 µm filter (CellTreat #229481), then again through a whatman grade 1 filter paper (Cytiva #1001-325) syringe filter (Fisher Scientific #NC9972954). Filtered cells were counted, then diluted to 2×10^5^ cells/mL into fresh 1% tryptone and seeded into a 12-well glass-like polymer bottom plate (Cellvis #P12-1.5P). The plate was sealed with parafilm and incubated at 24℃ for 48 hours. <u>Contractile vacuole and transition</u> <u>experiments</u>: *Bd* zoospores were harvested and filtered as for turgor pressure experiments. Filtered cells were centrifuged at 2500 rcf for five minutes, washed twice in 0.1X Bonner’s salts (1X Bonner’s salts:^59^ 10 mM sodium chloride, 10 mM potassium chloride, 2.7 mM calcium chloride), and resuspended in 2 mL 0.1X Bonner’s salts.

### Hyperosmotic shock of sporangia

<u>Short-term sorbitol treatments</u>: cells were stained for the cell wall using 0.1% calcofluor white with an Evans Blue counterstain (v/v; Sigma #18909) in 1% tryptone. After one minute of imaging, the media in the well was removed and replaced with either fresh 1% tryptone or 1% tryptone supplemented with 300 mM or 500 mM sorbitol (Sigma #S1876). The time of media replacement with respect to the start of the imaging acquisition was recorded. <u>Membrane and cell wall staining during hyperosmotic shock</u>: cells were incubated with 0.1% calcofluor white without any counterstain (v/v; Sigma #910090) for 20 minutes in 1% tryptone to stain the cell wall. Cells were washed three times with 1% tryptone and then stained for the cell membrane using 10 µM FM4-64 (Invitrogen #T13320). Sorbitol treatments were performed as in the short term experiments, except the treatment solution was supplemented with 10 µM FM4-64. <u>Long term sorbitol treatments</u>: cells were stained as in the short term experiments. After the first time lapse frame for each experimental condition was taken, fresh 1% tryptone or 1% tryptone supplemented with sorbitol was added to the appropriate wells to final concentration of 300 or 500 mM sorbitol. The second time lapse frame was considered to be the five minute post-treatment time.

### Contractile vacuole visualization in zoospores

<u>Sorbitol treatments under agarose</u>: 1.5% low-melt agarose (w/v; Thermo Scientific #R0801) pads were prepared using 0.1X Bonner’s supplemented with 0, 10, 25, 50, or 100 mM sorbitol. Concentrated cells prepared as described above (see “Cell Culture and Cell Preparation”) were added to the middle of a well of a 12-well glass-bottom plate (Cellvis #P12-1.5H-N). An agarose pad was gently placed over the cells and left for five minutes before excess liquid was removed from the edges of the pad via pipetting. To allow acclimation to the osmotic environment and adequate confinement, the cells were left for 30 minutes, during which any liquid around the edge of the pad was occasionally removed using the corner of a kim wipe. After this time, cells were imaged. <u>Membrane staining under</u> <u>agarose</u>: 1.5% low-melt agarose (w/v) pads supplemented with 10 µM FM4-64 were prepared using 0.1X Bonner’s salts. Concentrated cells were added to the center of a 12-well glass-bottom plate and FM4-64 was added to a final concentration of 10 µM. Then, the agarose pad was gently placed over the cells. After five minutes, excess liquid was removed via pipetting. After five more minutes, the remaining liquid around the edge of the agarose pad was removed with the corner of a kim wipe, and cells were imaged. <u>Chemical inhibitors under agarose</u>: 1.5% low-melt agarose (w/v) pads with either 0 or 50 nM Bafilomycin-A1 (Cayman Chemical #11038, resuspended in methanol) were prepared using 0.1X Bonner’s. Concentrated cells were added to the middle of a well of a 12-well glass-bottom plate and propidium iodide (PI, Invitrogen #P3566) was added to a final concentration of 0.1% (v/v). For treated cells, Bafilomycin-A1 was added to the well to a final concentration of 50 nM. The appropriate agarose pads were added over the cells and liquid was removed as described for the sorbitol treatments under agarose and cells were imaged.

### Encystation

Concentrated cells prepared as described above (see “Cell Culture and Cell Preparation”) were added to the middle of a well of a 6-well glass bottom plate (Cellvis #P06-1.5H-N) with calcofluor white with Evans Blue (Sigma #18909) at a final concentration of 0.1% (v/v). The solution was mixed gently before the addition of mucin (Sigma #M1778) to a final concentration of 10 mg/mL. Cells were then immediately confined using a dynamic cell confiner (4Dcell) with a PDMS suction cup and 1 µm pillar-height coverslip pre-soaked in 0.1X Bonners and 0.1% (v/v) calcofluor white solution. The coverslip was gently lowered onto the cells to prevent bursting and cells imaged after 15, 20, 25, 30, and 40 minutes of mucin treatment.

### Microscopy

All imaging was performed on an inverted microscope (Ti2-Eclipse; Nikon) at room temperature. OF-Brightfield, illumination supplied by a multi-wavelength LED lightsource (CoolLED pE-300 white); OF-Cy5, 550 nm illumination supplied by the included diodes in a multi-wavelength LED lightsource (CoolLED pE-300 white) at 5% power, and a multi-bandpass filter set (Chroma 89404) combined with a Chroma ET697/60m emission filter; OF-TRITC, same illumination supply, settings and multi-bandpass filter set as for OF-Cy5, but instead combined with a Chroma ET595/33m emission filter; HB-DAPI, 408 nm illumination at 25% power supplied by a Celesta Light Engine (Lumencor) with a Chroma ET460/50m emission filter; HB-FM4-64, 510 nm illumination at 50% power supplied by a Celesta Light Engine (Lumencor) with a Semrock 685/40 nm emission filter; AXR-FM4-64, 488 nm illumination supplied by a Nikon LUA-S4 laser launch at 50% power, and a proprietary quad-band emission filter (677/28); AXR-Trans, transmitted light simultaneously supplied by the 488 nm excitation light with a gain of 30 and pinhole size of 16.1. Zoospore imaging used a 1.5X tube lens in addition to the given objective described below.

<u>For short-term sorbitol treatments of sporangia</u>, cells were imaged with a 40x objective (plan flour, air, NA=0.60) using the OF-Cy5 configuration and a Prime BSI Express camera (Teledyne). Time lapse imaging was performed on the middle plane of the cells at 5 s intervals for 8 m with 400 ms exposure time. <u>For membrane and cell wall staining during hyperosmotic</u> <u>shock in sporangia</u>, cells were imaged with a 40x objective (plan apo λD air, NA=0.95) on a microscope equipped with a CrestV2 spinning disk confocal. The cell wall was visualized using the HB-DAPI configuration, the cell membrane was visualized with the HB-FM4-64 configuration, and all images were taken with a Prime 95B 25 mm camera (Teledyne). Time lapse imaging was performed with 200 ms exposure of 408 nm illumination followed by 300 ms exposure of 510 nm illumination at 10 s intervals for 8 m. <u>For long-term sorbitol treatments of</u> <u>sporangia</u>, cells were imaged with the same objective, configuration, and camera as for the short-term experiments. Time lapse imaging was performed on the middle plane of the cells at 5 m intervals for 120 m with 400 ms exposure time. Three XY points per well were chosen for each treatment and all XY points were acquired in the same imaging acquisition using Nikon Elements v6.02.03.

<u>For sorbitol and Bafilomycin A1 under agarose experiments</u>, cells were imaged with a 100x objective (plan apo λ oil, NA=1.45) using the OF-Brightfield configuration and a Prime BSI Express camera (Teledyne); the OF-TRITC configuration was also used when applicable to visualize PI staining. Timelapse images were taken with 50 ms brightfield exposure (iris open 3.3% or 5%), followed by 100 ms of 550 nm excitation when applicable, every 250 ms for 3 m. <u>For under agarose membrane staining in zoospores</u>, cells were imaged on a microscope equipped with an AX R point scanning confocal system with a Nikon Spatial Array Confocal detector. Cells were imaged with a 60x oil objective (plan apo λD oil, NA=1.42) using the AXR-Trans and AXR-FM4-64 configurations. Images were taken at 5 s intervals for 2 m using a galvano unidirectional band scanner with a dwell time of 0.4 µs. <u>For the mucin-induced</u> <u>encystation experiments</u>, cells were imaged with a 100x objective (plan apo λ oil, NA=1.45) using the OF-Brightfield configuration, OF-Cy5 configuration, and a Prime BSI Express camera (Teledyne) to visualize whole cells and the cell wall via Evans Blue. Timelapses were taken at 250 ms intervals for 1.5 m using 50 ms brightfield exposure (iris open 3.3%) and 50 ms 500 nm illumination exposure every 100 frames.

### Quantification and Statistical Analysis

All experiments were performed on three independent populations of sporangia or zoospores harvested on different days (biological replicates).

<u>For short-term time lapse of sporangia</u> (**Figure 1A-B**), the frames corresponding to one minute before and one, three and five minutes after media replacement were isolated and then processed using a custom General Analysis 3 pipeline in Nikon Elements v6.02.03 (**File S1**). Each image was adjusted to increase contrast using a rolling ball average, then thresholded to identify individual sporangia over time. Objects with diameters less than 10 µm, along with those that touched the edges of the image, were discarded, and the area of remaining objects measured. Prior to any calculations, objects not present in all four frames as well as objects with areas below 130 µm^2^ or above 500 µm^2^ (objects representing partial or multiple adjacent cells) were removed using a custom python script (**File S2**). The percent change in cross-sectional area for each cell was then calculated by dividing the area of each cell in each after-treatment frame by the area of the same cell in the before-treatment frame. Statistical analysis between each treatment was performed on the average percent change in area for each of the three biological replicates using a one-way ANOVA with Tukey’s multiple comparisons. <u>To quantify</u> <u>plasma membrane delamination</u> (**Figure 1C-D**), samples were blinded and delamination was manually quantified as a clear separation between the plasma membrane and cell wall signal, or plasma membrane circles adjacent to the cell periphery. A one-way ANOVA with Tukey’s multiple comparisons on the replicate averages for each sorbitol treatment was performed. <u>For</u> <u>long-term time lapse of sporangia</u> (**Figure 1E**), individual XY points were processed using the same General Analysis 3 pipeline used for the short term experiments (**File S1;** binary shown in **Figure S1A**; example cells shown in **Figure S1B**), objects not in all 25 time frames removed, and the same area cut-off range filter used as for the short term experiments. For all 25 frames, the percent change in area for each cell was calculated as for the short term experiments using a custom python script (**File S3**).

<u>For sorbitol treatments under agarose</u> (**Figure 3A**), samples were blinded, five representative cells were selected per treatment condition, and image frames corresponding to the first 60 seconds of imaging were analyzed using Fiji. (v1.53r).^60^ Each vacuole that disappeared was counted towards the pumping rate. The area of each shrinking vacuole was measured at its largest using the wand (tracing) tool or by freehand selection tool as was the area of each cell. The “percent cell area pumped per minute” was determined by dividing the sum of contractile vacuole areas at their largest by the cell’s area. To quantify contractile vacuole dynamics over time (**Figure 2A**), a representative cell from the 0 mM sorbitol condition was chosen, and every contractile vacuole was measured for each frame of the video for 60 seconds. <u>For Bafilomycin-</u> <u>A1 treatments under agarose</u> (**Figure 3B**), contractile vacuole sizes, pumping rate, and percent of cell area pumped per minute were determined using the same methods as for the under agarose sorbitol treatments. Cell death (**Figure S4C**) was evaluated by positive PI staining. Cells that were clearly encysted (small and nearly perfect circles), that touched the border of the image, or were out of focus were not included in analysis. Two-tailed Student’s t-tests were performed on the measurements for multiple pumps and cell death. <u>For encystation</u> <u>experiments</u> (**Figure 4**), time points were blinded, five randomly selected cells per replicate chosen, and the average cell wall intensity for the first frame was measured using the free-hand selection tool in Fiji to outline the cell. The pumping rate at each time point was quantified as described for the sorbitol treatments under agarose. The time between appreciable cell wall accumulation and the cessation of vacuole pumping was defined as the time difference between the first time point at which the cell’s total cell wall intensity was at least 20% brighter than background and the time point in which a cell had no pumping contractile vacuoles.

### High Osmolarity Glycerol Pathway Homolog Identification

Putative homologs of key components of the HOG pathway in *Bd* and other chytrids were identified by ortholog group analysis, domain analysis, and database and literature searches (**Figure S2** and **Data S2A**). The following species were included in this analysis: *Allomyces macrogynus* ATCC38327 (*Am*; GenBank: GCA_000151295.1; WGS project: PRJNA20563); *Aspergillus nidulans* FGSC A4^61^ (*An*; NCBI RefSeq: GCF_000011425.1); *Batrachochytrium dendrobatidis* JEL423^62^ (*Bd*; GenBank: GCA_000149865.1); *Candida albicans* SC5314^63^ (*Ca*; NCBI RefSeq: GCF_000182965.3); *Magnaporthe (Pyricularia) oryzae* 70-15^64^ (*Mo*; NCBI RefSeq: GCF_000002495.2); *Neurospora crassa* OR74A^65^ (*Nc*; NCBI RefSeq: GCF_000182925.2); Saccharomyces *cerevisiae* S288C^66^ (*Sc*; NCBI RefSeq: GCF_000146045.2); *Schizosaccharomyces pombe* 972h-^67^ (*Spo*; NCBI RefSeq: GCF_000002945.1); and *Spizellomyces punctatus* DAOM BR117^68^ (*Sp*; NCBI RefSeq: GCF_000182565.1). For all proteins, splice variants were not considered.

HOG pathway mitogen activated kinase (MAPK) cascade components were collected from Table S4 of a previous study.^69^ For each protein and species of interest, the reported GenInfo Identifiers (gi) were used to identify the associated NCBI/GenBank entry and accession number. In any cases where the gi linked to non RefSeq assemblies, or records that were superseded or suppressed, BLAST^70^ was used to identify the protein in the RefSeq assembly.

Non-MAPK cascade proteins in the HOG pathway were identified by searching for *Am*, *Bd*, and *Sp* orthologs in OrthoMCL database^71,72^ orthogroups that contained known S. *cerevisiae (Sc)* HOG pathway proteins (**Data S1A**). The protein domains of the resulting ortholog candidates were analyzed using InterProScan with default settings^73^ and chytrid proteins missing one or more domains predicted to be in the *Sc* protein were removed from the dataset. If an orthogroup did not produce a candidate protein in a given species, BLASTp and/or tBLASTn (via NCBI) was used to manually identify a potential candidate using the *Sc* protein sequence as the query. For BLASTp, the nr database was used, filtering results to consider only the chytrid species of interest, a word size of three, and all other parameters set to the default settings. For tBLASTn, the search was performed on default settings against only the species of interest’s genome. The top five hits from each species were then used as the queries to do a reverse BLASTp search using the RefSeq database and filtering to consider only *Sc* (parameters as described for forward BLASTp search). Chytrid proteins for which their mutual best BLAST hit (MBBH) was the query *Sc* protein, and also contained all the expected domains from InterProScan, were considered a putative homolog. If still no clear homolog was found, then the protein was marked as not found.

Identification of non-MAPK cascade proteins in the HOG pathway of Dikarya (*An*, *Ca*, *Mo*, *Nc*, and *Spo*): *Spo*, the “protein with orthologs” function on pombase.org^74^ was used to identify proteins that have an ortholog in *Sc*, followed by finding each protein’s corresponding NCBI RefSeq accession number; *Ca*, each *Sc* protein was searched for on candidagenome.org,^75^ ensuring that the *Sc* protein was under the “ortholog(s) in non-CGD species’’ section in the given gene page and the “external links” section was used to find the corresponding NCBI RefSeq accession; *An*, *Mo*, and *Nc* the “identify genes based on a list of IDs” option on fungidb.org^76,77^ was used. The list of *Sc* systematic names was used as the input and a step was added to transform the results into orthologs in each of the species of interest. Entrez and Uniprot IDs for the predicted orthologs were downloaded and the corresponding NCBI RefSeq accession numbers were identified. If no ortholog for a protein in a Dikarya species was found through these methods, the MBBH hit approach outlined above was used, using the RefSeq database and filtering for the species of interest only for the forward search. If still no ortholog for a protein was found, a literature review for the protein of interest was performed. The protein was marked as not found if none of the above methods returned a candidate.

### Vacuolar ATPase and Aquaporin Homolog Identification

Domain analysis and BLAST searches were performed to identify homologs of v-ATPase subunits and aquaporins (**Figure S3** and **Data S2B**) in the same species of interest as with the HOG pathway homolog search, with these additional species: *Acanthamoeba castellanii* str. Neff^78^ (*Ac*; RefSeq: GCF_000313135.1); *Arabidopsis thaliana*^79^ (*At*; RefSeq: GCF_000001735.4); *Chlamydomonas reinhardtii*^80^ (*Cr*; RefSeq: GCF_000002595.2); *Dictyostelium discoideum* AX4^81^ (*Dd*; RefSeq: GCF_000004695.1); *Homo sapiens* (*Hs*; RefSeq: GCF_000001405.40); *Naegleria gruberi*^82^ (*Ng*; RefSeq: GCF_000004985.1); *Trypanosoma brucei* 927/4 GUTat10.1^83^ (*Tb*; RefSeq: GCF_000002445.2); and *Trypanosoma cruzi* CL Brener^84^ (*Tc*; RefSeq: GCF_000209065.1). Splice variants were not considered.

The v-ATPase is a multisubunit protein complex.^48^ For *Sc*, each subunit of the v-ATPase can be uniquely identified via Interpro family or domain membership. The most descriptive Interpro entry accession numbers for each subunit from *Sc* were identified (**Data S1B**). The *c*, *c’*, and *c”* subunits are difficult to distinguish from each other,^49^ so were combined into one category. Homologs were identified by inclusion in each subunit’s Interpro entry and their associated NCBI/GenBank accession numbers identified through their Uniprot entry page or BLAST via NCBI. If no candidate subunit homologs were identified in a given species, manual BLAST searches were performed as for the HOG pathway proteins. If a protein could still not be identified, it was marked as not present in the species. The *c*/*c’*/*c”* and *a* subunits from *Bd*, *Sc*, and *Dd* were aligned (**Figure S4A**) using TCoffee on default settings.^85^

Aquaporins were identified by inclusion in the Interpro major intrinsic protein (MIP, IPR000425) family.^86,87^ When available, only reviewed protein entries were counted. All proteins’ associated NCBI/GenBank accession numbers were found through their Uniprot entry page or BLAST via NCBI. Inactive accession numbers were removed from the dataset.

---

**Figure S1.**
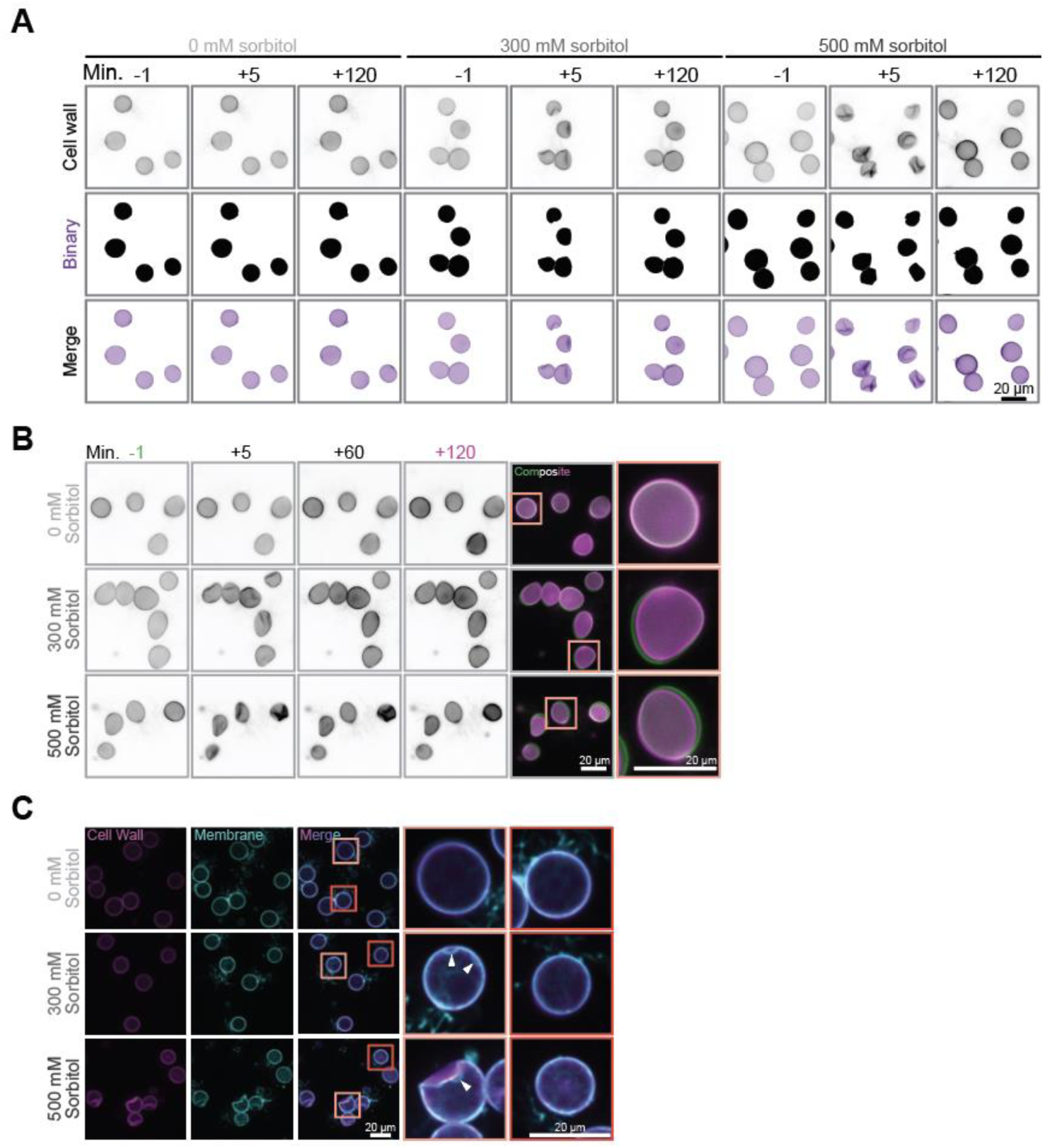
Analysis of *Bd* sporangia responding to hyperosmotic shock. **(A)** Representative images of *Bd* sporangia cell walls stained with Evans Blue (top) one minute before (−1) and five (+5) and 120 (+120) minutes after treatment with media supplemented with the indicated sorbitol concentration. All cell wall images are adjusted to the same brightness and contrast. Cells were segmented using the cell wall signal in NIS elements (v6.02.03), resulting in a binary layer (purple) that encompasses the cell body **(B)** Representative images of *Bd* sporangia cell walls stained with Evans Blue one minute before (−1) and five (+5), 60 (+60) and 120 (+120) minutes after treatment with media supplemented with the indicated concentration of sorbitol. Composite images show cells one minute before (green) and 120 minutes after (magenta) sorbitol treatment. All images are adjusted to the same brightness and contrast. **(C)** Example images of *Bd* sporangia stained for the cell wall (magenta) and membrane (cyan) after five minutes of treatment with media supplemented with the given sorbitol concentration. Merged images show the overlay of membrane and cell wall. For the 300 and 500 mM sorbitol treatments, insets give one example of a cell with delamination and another example of a cell without delamination from the same field of view. White arrowheads indicate membrane delaminations. All images for each stain are adjusted to the same brightness and contrast.

**Figure S2.**
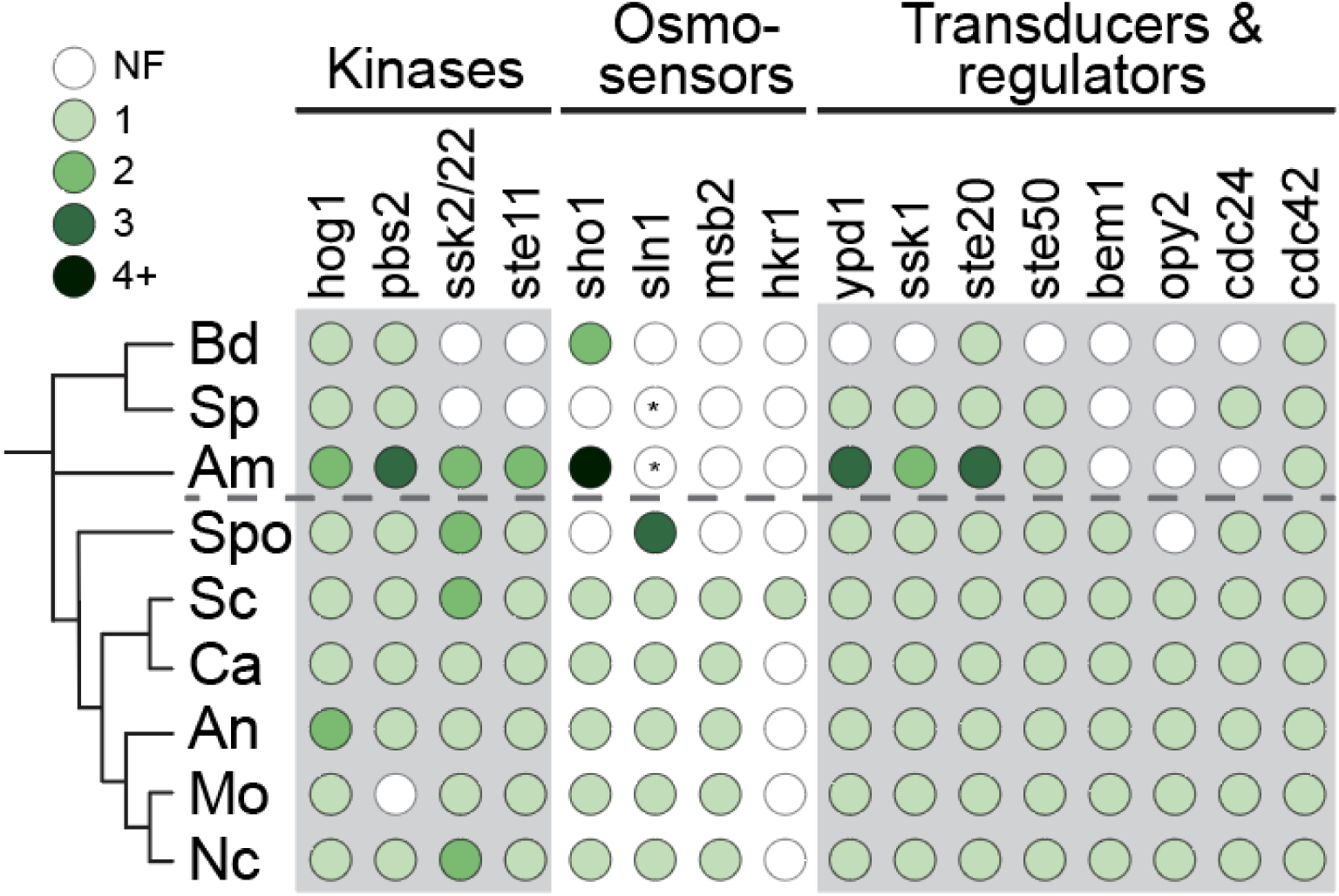
*Bd* and other chytrids have putative homologs of several key components of the HOG pathway. The distribution of proteins known to function in the high osmolarity glycerol pathway (HOG) across fungal taxa. The HOG pathway is the primary pathway used by budding yeast to regulate turgor pressure and has three main classes of proteins: 1) a mitogen activated kinase (MAPK) cascade ending with the MAPK Hog1; 2) osmo-sensors to sense changes in external osmolarity; and 3) transducers and regulators that connect the MAPKs and osmosensors. White-filled circles indicate that homologs are not found (NF), color-filled circles indicate the detection of one or more homologs. Kinase copy numbers were obtained from.^69^ Dashed line separates chytrid species (above) from Dikaryotic species (below). *: There is no clear homolog for sln1, but there are putative histidine kinases with predicted transmembrane domains in the given species’ genomes. Transmembrane domains are a hallmark of sln1-related histidine kinases.^88,89^ *Am*, *Allomyces macrogynus*; *An*, *Aspergillus nidulans*; *Bd*, *Batrachochytrium dendrobatidis*; *Ca*, *Candida albicans*; *Mo*, *Magnaporthe oryzae*; *Nc*, *Neurospora crassa*; *Sc*, *Saccharomyces cerevisiae*; *Sp*, *Spizellomyces punctatus*; *Spo*, *Schizosaccharomyces pombe*.

**Figure S3.**
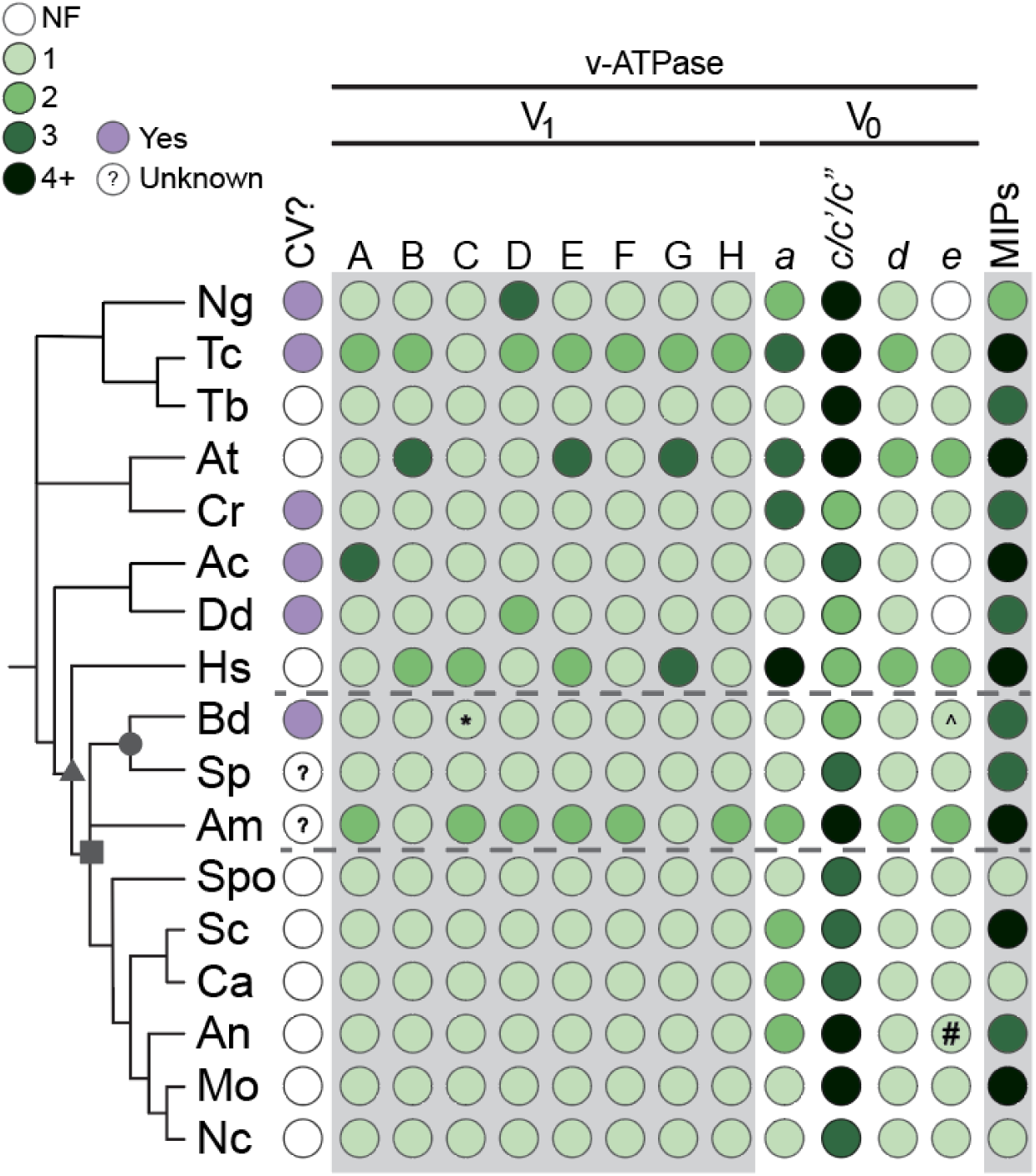
*Bd* and other chytrids have putative aquaporin and vacuolar-ATPase homologs. The distribution of major intrinsic family proteins (MIPs), subunits of the vacuolar-ATPase (v-ATPase), and presence of contractile vacuoles (CV) across taxa. Aquaporins are part of the MIP family of proteins. The v-ATPase is made up of two complexes (V_1_ and V_0_), each comprising several subunits.^48^ White-filled circles indicate that homologs are not found (NF), color-filled circles indicate the detection of one or more homologs. Purple circles indicate the presence of documented CVs in the given organism. *: Homolog is predicted to be in the ubiquitin activating enzyme family (IPR018075), but this likely represents two separate genes erroneously annotated as one. ^: Homolog only identifiable when using the *Manduca sexta* protein (NCBI RefSeq XP_037299296.1) as a query for BLASTp or tBLASTn. #: Homolog is not annotated in the reference genome, and is only identifiable when using the *Sc* protein as a query for tBLASTn. Symbols on the tree represent opisthokonts (triangle), fungi (square), and Chytridiomycota (circle). Dashed lines surround chytrid species. *Ac*, *Acanthamoeba castellanii*; *Am*, *Allomyces macrogynus*; *An*, *Aspergillus nidulans*; *At*, *Arabidopsis thaliana*; *Bd*, *Batrachochytrium dendrobatidis*; *Ca*, *Candida albicans*; *Cr*, *Chalmydomonas reinhardtii*; *Dd*, *Dictyostelium discoideum*; *Hs*, *Homo sapiens*; *Mo*, *Magnaporthe oryzae*; *Nc*, *Neurospora crassa*; *Ng*, *Naegleria gruberi*; *Sc*, *Saccharomyces cerevisiae*; *Sp*, *Spizellomyces punctatus*; *Spo*, *Schizosaccharomyces pombe*; *Tb*, *Trypanosoma brucei*; *Tc*, *Trypanosoma cruzi*.

**Figure S4.**
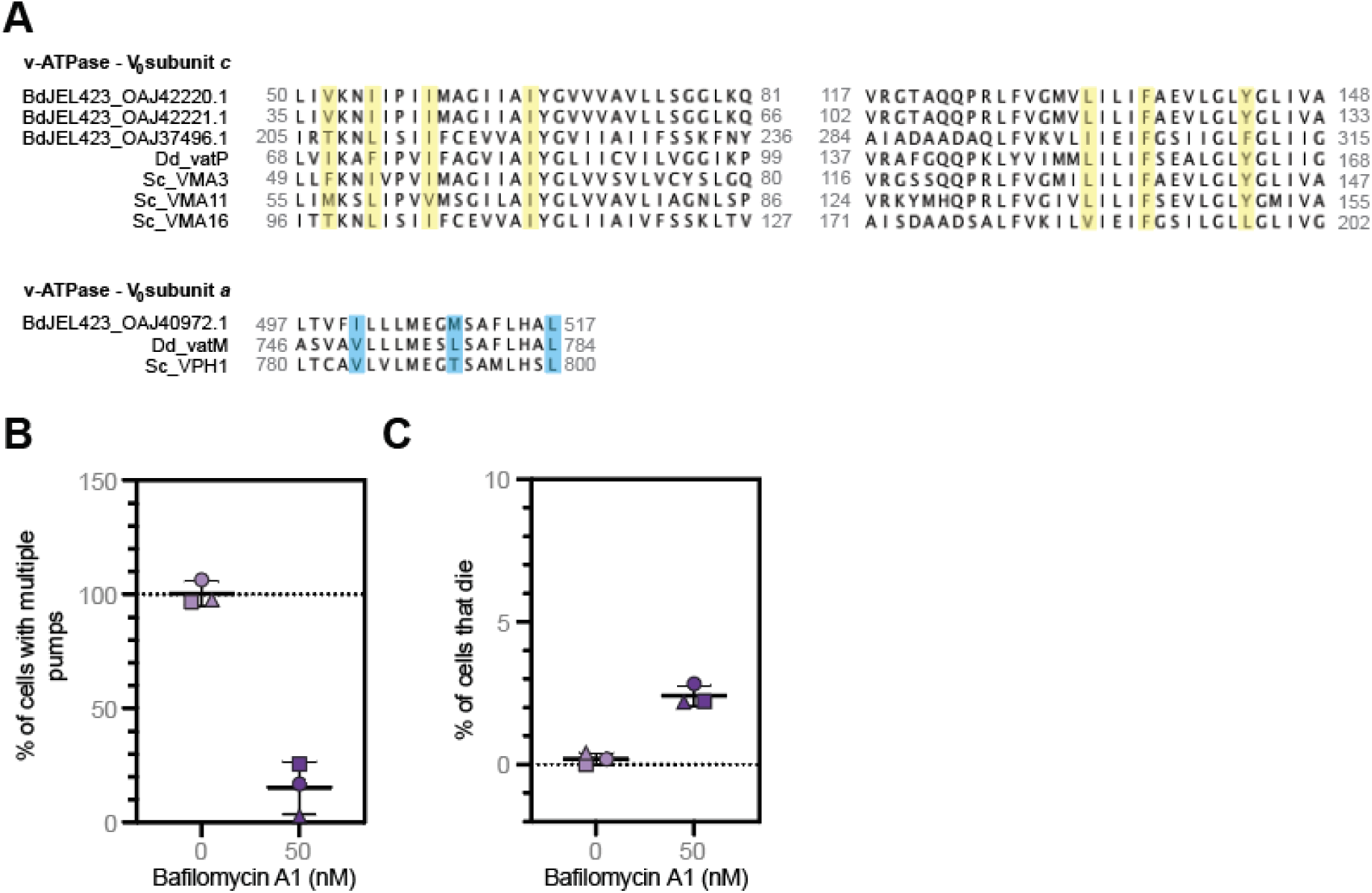
*Bd* is susceptible to the vacuolar ATPase inhibitor Bafilomycin A1. (**A**) TCoffee alignment of known and putative homologs for the indicated vacuolar ATPase subunits in the given species. Residues that form the bafilomycin A1 binding site (yellow) or that are disrupted by bafilomycin A1 binding (cyan) are highlighted. *Bd*, *Batrachochytrium dendrobatidis* strain JEL423; *Dd*, *Dictyostelium discoideum* AX4; *Sc*, *Saccharomyces cerevisiae* 288C. (**B**) Quantification of the percent of *Bd* zoospores that exhibit organelles undergoing multiple growing and shrinking cycles over a three minute period under agarose treated with 0 or 50 nM bafilomycin A1. Three independent biological replicates were performed, each represented by a shape. Mean and standard deviation of the three biological replicates are indicated by black lines. Two-tailed Student’s t-test: p = 0.0007. **(C)** Quantification of the percent of *Bd* zoospores that die over a three minute period under agarose treated with 0 or 50 nM bafilomycin A1. Calculated by taking the difference between the percent of propidium iodide positive cells between the last and first frames of the time lapse. Three independent biological replicates were performed, each represented by a shape. Mean and standard deviation of the three biological replicates are indicated by black lines. Two-tailed Student’s t-test: p = 0.0008.

## LEGENDS FOR SUPPLEMENTAL MATERIAL

**Supplemental Data 1. Identification numbers used in determining high osmolarity glycerol pathway and vacuolar ATPase homologs. (A)** Orthogroup numbers from the OrthoMCL database to which proteins from the high osmolarity glycerol pathway in *Saccharomyces cerevisiae* belong. These ortholog groups were used in this study to determine putative orthologs for the given protein in chytrid species. **(B)** Interpro family or domain accession numbers that uniquely identify *Saccharomyces cerevisiae* vacuolar ATPase subunits. These Interpro accession numbers were used to identify vacuolar ATPase subunits in chytrid species by searching the proteins classified under each entry.

**Supplemental Data 2. Accession numbers for putative homologs of proteins involved in osmoregulation in the given species. (A)** NCBI/GenBank accession numbers for putative homologs of proteins involved in the high osmolarity glycerol pathway. The mitogen activated protein kinase family members were identified based on a previous study which identified these proteins in the given species based on phylogenetic analysis.^69^ **(B)** NCBI/GenBank accession numbers for putative homologs of vacuolar ATPase subunits and members of the major intrinsic protein super family (includes aquaporins).

**Supplemental File 1. Area thresholding analysis pipeline for sorbitol treatments on *Batrachochytrium dendrobatidis* sporangia.** Nikon Elements General Analysis 3 pipeline for thresholding cells to determine cross sectional area of *Batrachochytrium dendrobatidis* sporangia before and after sorbitol treatment. Pipeline tracks each cell through time and thresholds based on cell wall signal to measure diameter and area of the cell. For the analysis in this study, the rolling ball average was set to 100% degree and 0.49 µm radius. The threshold used was 450-inf, with smooth set to 0.41 µm, clean set to 0.25 µm, separate set to 0.082 µm and fill holes option on. A size filter of 10-1000 µm was also applied. This file was used for both the short-term and long-term sorbitol treatments.

**Supplemental File 2. Python script for processing area measurements for short-term sorbitol treatments on *Batrachochytrium dendrobatidis* sporangia.** Custom python script written to process the thresholding measurements for *Batrachochytrium dendrobatidis* for short term sorbitol treatment experiments. Data input ideally comes from a Nikon Elements General Analysis 3 pipeline that tracks objects through time. This script filters out thresholded object groups (one group is a cell over time) that are not in all of the given number of frames, as well as object groups where at least one of the area measurements for a time frame is outside of a given range. Then, this script will determine the percent change in area and diameter for each cell based on the cell’s area and diameter in the first time frame. Output will be the filtered original data, the percent change in area and diameter for each time point (e.x., 1 minute before, and 1, 3, and 5 minutes after treatment) as well as averages for all cells per treatment per replicate. Output is formatted for easy input into GraphPad Prism software. For this study, the number of time frames was set to four; the lower area threshold was 130 and the upper area threshold was 500.

**Supplemental File 3. Python script for processing area measurements for long-term sorbitol treatments on *Batrachochytrium dendrobatidis* sporangia.** Custom python script written to process the thresholding measurements for *Batrachochytrium dendrobatidis* for long term sorbitol treatment experiments. Data input ideally comes from a Nikon Elements General Analysis 3 pipeline that tracks objects through time. This script filters out thresholded object groups (one group is a cell over time) that are not in all of the given number of frames, as well as object groups where at least one of the area measurements for a time frame is outside of a given range. Then, this script will determine the percent change in area and diameter for each cell based on the cell’s area and diameter in the first time frame. Output will be the filtered original data, the percent change in area and diameter for each time point (e.x., 1 minute before, and 5, 10, 15 minutes after treatment) as well as averages and standard deviations for all cells per treatment per replicate. Output is formatted for easy input into GraphPad Prism software. For this study, the number of time frames was set to 25; the lower area threshold was 130 and the upper area threshold was 500.

**Supplemental Video 1. *Bd* sporangia exhibit membrane delamination upon hyperosmotic shock.** Representative time lapses of *Bd* sporangia stained for the cell wall (magenta) and membrane (cyan) throughout treatment with media supplemented with the given sorbitol concentration. Merged images show the overlay of membrane and cell wall. Cells are the same as shown in Figure 1C.

**Supplemental Video 2. *Bd* sporangia recover volume after hyperosmotic shock.** Representative time lapses of *Bd* sporangia stained for the cell wall (black) throughout 120 minutes of treatment with media supplemented with the given sorbitol concentration. Cells are the same as shown in Figure S1B.

**Supplemental Video 3. *Bd* zoospores have small organelles that grow and shrink over time.** Representative brightfield time lapse of *Bd* zoospores under agarose in 0.1X Bonner’s salts. Arrows and arrowheads indicate example organelles. Brightfield is on an inverted LUT. Cell is the same as shown in Figure 2A.

**Supplemental Video 4. *Bd* zoospores’ growing and shrinking organelles are membrane-bound.** Representative time lapse of *Bd* zoospores stained for membrane (black or cyan) under agarose in 0.1X Bonner’s salts. Merged images show the overlay of brightfield and membrane staining. Arrowheads indicate example organelles. Brightfield is on an inverted LUT. Cell is the same as shown in Figure 2B.

**Supplemental Video 5. Small, membrane-bound organelles in *Bd* zoospores respond to hyperosmotic shock and rely on vacuolar-ATPase function.** (Top) representative brightfield time lapse of *Bd* zoospores under agarose in 0.1X Bonner’s salts supplemented with the given sorbitol concentration. (Bottom) representative brightfield time lapse of *Bd* zoospores under agarose in 0.1X Bonner’s salts treated with the given concentration of the vacuolar ATPase inhibitor Bafilomycin A1. All brightfield images are on an inverted LUT. Arrows indicate example contractile vacuoles. Cells are the same as shown in Figure 3.

## Notes

### Competing Interest Statement

The authors have declared no competing interest.

