## Supplemental files 1-3 for "Frog-killing chytrid fungi deploy different strategies to regulate intracellular pressure in cell types that have or lack a cell wall": 2025_Prostak_Osmoregulation_File_S2.py

```
## will take ga3 data for turgor sorbitol experiments
## will calculate the % change in EqDia and Area across each frame in comparison to the before treatment frame
## will format the data for easy input into Prism

## input document must be a .csv with the following columns:
##	FileName	TrackId	Replicate	Treatment[mM]	TimeLapseIndex	ObjectId	BinObjectArea[um2]	BinEqDiameter[um]	BinCircularity

## FileName must NOT have spaces

# import numpy as np 
import pandas as pd
from datetime import datetime


def get_rep_avgs(filtered_df, list_to_groupBy, columns_to_avg):
	'''
	Calculates the average of desired columns in a dataframe based on a grouping, like replicate and treatment
	
	Parameters:
		filtered_df	(pd.DataFrame): the dataframe of objects after filtering out non-analyzable objects
		list_to_groupBy (list): list of column names in the dataframe to group rows by, for example:
			("ReplicateID", "Treatment")
		columns_to_avg (list): list of column names in the dataframe to be averaged per group, for eaxmple:
			("ratio_1-1", "ratio_2-1", "ratio_3-1", "ratio_4-1")
			
	Returns:
		repLevel_avgs (pd.DataFrame): a dataframe of the averages for each group for each column desired
	'''
	
	repLevel_avgs = filtered_df.groupby(list_to_groupBy)[columns_to_avg].mean().reset_index()
	print(f'''The rep avgs are: \n{repLevel_avgs}\n\n\n''')
	return repLevel_avgs

def get_cellLevel_diffs(filtered_df, measurement, columns_to_keep):
	'''
	NOTE: this function is not used in version 2.1, the input ga3 data does not support this,
		and difference are not of interest anymore, only percents 
	
	Calculates the difference between a given measurement before and after treatment and removes unwanted columns
	
	Parameters:
		filtered_df	(pd.DataFrame): the dataframe of objects after filtering out non-analyzable objects
		measurement (str): the column name of the measurement to calculate
		columns_to_keep (list): list of columns to keep in the output dataframe
	
	Returns:
		cellLevel_df (pd.DataFrame): a dataframe with the differences for each cell
	'''

	cellLevel_df = filtered_df.dropna(subset=[measurement])[columns_to_keep]
# 	print(f'''The raw cell level data are: {cellLevel_df}''')
	return cellLevel_df

def combine_values(row):
	'''
	Combines values for a cell to create a unique identification string for ease of reference and data handling
	
	Parameters:
		row (pd.DataFrame): a row of the filtered dataframe of interest
		
	Returns:
		 a string uniquely representing the cell in the given row
	'''
	
	date = row['FileName'].split('_')[0]
	treatment = str(row['Treatment[mM]']).split(".")[0]
	file_num = row['FileName'].split('_')[4]
	rep = row['Replicate']
	cell_num = row['TrackId']
	return f'''{date}_{treatment}_{file_num}_{rep}_{cell_num}'''

def collapse_cellID_info(cellLevel_df, new_column_name, columns_to_remove, new_column_order):
	'''
	Adds a cell's unique identification string to a new column, removes unwanted columns, and reorders columns as desired.

	Parameters:
		cellLevel_df (pd.DataFrame): the dataframe of objects after filtering out non-analyzable objects
		new_column_name (str): the name of the column of ID strings
		columns_to_remove (list): a list of column names of columns to be removed
		new_column_order (list): a list of column names in the desired order
		
	Returns:
		cellLevel_df (pd.DataFrame): the input dataframe of objects, with the new column added, 
			unwanted columns removed, and columns in the desired order
	'''

	cellLevel_df[new_column_name] = cellLevel_df.apply(combine_values, axis=1)
	cellLevel_df = cellLevel_df.drop(columns=columns_to_remove)
	cellLevel_df = cellLevel_df[new_column_order]
# 	print(f'''\n\nThe cell level data with unique cellIDs are:\n{cellLevel_df}''')
	return cellLevel_df

def pivot_df(collapsed_cellLevel_df, index_to_pivot, column_to_pivot, measure):
	'''
	Pivots a given dataframe for proper output into a .csv for a desired measure
	
	For example: 
	
		ID	treatment	ratio_1-1	ratio_2-1
		A	0			1			0.95
		B	300			1			0.5
		
	If index_to_pivot = "ID", column_to_pivot= "treatment", and measure= "ratio_2-1",
		the pivoted table will be:
		
		ID	0	300
		A	1	
		B	nan	0.5
		
		
	Parameters:
		collapsed_cellLevel_df (pd.DataFrame): a dataframe of measurements for each cell identified by their unique string
		index_to_pivot (str): the name of the column to pivot to a row of the new dataframe
		column_to_pivot (str): the name of the column to pivot to a column of the new dataframe
		measure (str): the name of the column where the values will be pivoted to the values of the new dataframe
	
	Returns:
		pivoted_df (pd.DataFrame): the dataframe with the desired pivot
	'''

	pivoted_df = collapsed_cellLevel_df.pivot(index=index_to_pivot, columns=column_to_pivot, values=measure).reset_index()
	pivoted_df.columns.name = None
# 	print(f'''The pivoted cell level data are:\n{pivoted_df}''')
	return pivoted_df
	
def save_to_csv(dataframe, output_directory, measure, level):
	'''
	Saves a dataframe as a .csv to a given directory, saving the file name using the inputed variables
	
	Parameters:
		dataframe (pd.DataFrame): the dataframe to be saved
		output_directory (str): the path to the directory in which the file is to be saved
			must NOT have ending /
		measure (str): the measure being saved (ex., "ratio2-1")
		level (str): the level of the measurement being saved (i.e., cell level, replicate average)
		
	Returns:
		None, writes a .csv to indicated directory
	'''
	
	file_path = f'''{output_directory}/{current_date}_{measure}_{level}.csv'''
	dataframe.to_csv(file_path, index=False)
	print(f'''The output for {measure}, {level} has been written to:\n{file_path}\n\n\n''')
	return None

def calculate_cellLevel_pcts(group, column):
	'''
	Calculates the percent change in a given column before and after treatment
	Determines the reference value for the column as the value with a TimeLapseIndex of 1
	
	Use within the .apply method for a grouped dataframe
	
	Parameters:
		 group (grouped pd.DataFrame): the filtered dataframe with unique cell ID strings, 
		 	grouped according to cell and treatment
		 column (str): the name of the column to be used to calculate the percent changes (ex. "Area")

	Returns:
		group (grouped pd.DataFrame): the grouped dataframe, with the calculated percent changes added as new columns
	'''

	reference_diameter = group.loc[group['TimeLapseIndex'] == 1, column].values[0]
	for TLindex in [1, 2, 3, 4]:
		group[f'Ratio_{TLindex}-1'] = group.loc[group['TimeLapseIndex'] == TLindex, column].values[0] / reference_diameter * 100
	return group


##########################################################################################
######################### BEGIN USER DEFINED INPUT #######################################

## No change needed: sets current date of when the script is run
current_date = datetime.now().strftime('%Y-%m-%d')

## the number of frames an object needs to be present for analysis
frame_num = 4

## the area on the upper limit of the distribution to remove objects that are likely multiple cells
## in um^2, as an integer
upper_area_cutoff = 500

## the area on the lower limit of the distribution to remove objects that are likely improperly recognized cells
## in um^2, as an integer
lower_area_cutoff = 130

## Path to input .ga3 results file
ga3_file = "/Users/laboratory/Documents/Fritz_Laylin_Lab/Bd_osmoreg/turgor/Sorb/GA3_output/GA3_v8_output_run_11-12-24_analyzable.csv"

## Path to DIRECTORY to save output files to-- 
## individual files will be saved with unique file names according to 
## current date, measurement type, and cell vs replicate level
## DO NOT include the final "/" in the directory
output_directory = '/Users/laboratory/Documents/Fritz_Laylin_Lab/Bd_osmoreg/turgor/Sorb/replicates_for_prism/short-term_redo/with_area'


#----------------------------------------------------------------------------#									
## The following lists must match the names of the columns in the .ga3 file ##
#----------------------------------------------------------------------------#

## List of columns to keep when filtering out cells with data for fewer than 4 frames
selected_columns = ['FileName', 'TrackId', 'Replicate', 'Treatment[mM]', 'BinObjectArea[um2]', 
					'BinEqDiameter[um]']

EqDia_columnName = 'BinEqDiameter[um]'

Area_columnName = 'BinObjectArea[um2]'

## List of columns to remove after creating a unique cellID column for pct change calculation
columns_to_remove_pct = ['FileName', 'TrackId', 'Replicate', 'ObjectId', 'BinCircularity']

#----------------------------------------------------------------------------#									
##---------------- lists for the percent change values ---------------------##
#----------------------------------------------------------------------------#


## List of column names for the percent change in area columns to be used in pivoting						
cellLevel_pcts = ['Ratio_1-1', 'Ratio_2-1', 'Ratio_3-1', 'Ratio_4-1']

## List of column names to keep when trimming down the cell-level dataframe
columns_to_extract = ['UniqueCellID', 'Ratio_1-1', 'Ratio_2-1', 'Ratio_3-1', 'Ratio_4-1', 'Treatment[mM]']

## List of column names to group on when averaging percent changes per replicate
avging_groups_pcts = ['GroupID', 'Treatment[mM]']


######################### END USER DEFINED INPUT #########################################
##########################################################################################


## import the ga3 data into a pandas dataframe
og_df = pd.read_csv(ga3_file)
# print(og_df)


## group the data by file name and trackID so all four frames for one cell stay together 
grouped_df = og_df.groupby(['FileName', 'TrackId'])
# print(grouped_df)

# for key, item in grouped_df:
#     print(grouped_df.get_group(key), "\n\n")

# Filter out cells that do not fit the frame number and area criteria for analysis
filtered_df = grouped_df.filter(
    lambda group: (
        group['TimeLapseIndex'].count() == frame_num and  # Ensure all frames are present
        all((lower_area_cutoff <= group['BinObjectArea[um2]']) & (group['BinObjectArea[um2]'] <= upper_area_cutoff)) # Exclude groups where at least one value is outside of the cutoff range
    )
)

print(filtered_df)


## save the filtered data before doing any further dataframe manipulation
save_to_csv(filtered_df, output_directory, 'filtered', 'data')


#----------------------------------------------------------------------------#									
##----------------- processing the percent change values -------------------##
#----------------------------------------------------------------------------#

## determine the new column order
pct_column_order = ['UniqueCellID'] + list(filtered_df.columns.difference(columns_to_remove_pct))

## create and add the unique identification strings for the cells, remove unwanted columns, reorder columns
uniqueID_df = collapse_cellID_info(filtered_df, 'UniqueCellID', columns_to_remove_pct, pct_column_order)
# print(uniqueID_df)


##the below commented out block of code is to check to see what groups may be missing a TLIndex equal to 1
## I put this in here as a test in case I missed a row when adding treatment values for the input
# test_df = uniqueID_df.groupby(['UniqueCellID', 'Treatment[mM]'], group_keys=False)
# print(f'''\n\ntest df: {test_df}\n\n''')
# missing_tlindex_groups = []
# 
# # Iterate through each group
# for name, group in test_df:
# 	# Check if there is any row where TimeLapseIndex == 1
# 	if not (group['TimeLapseIndex'] == 1).any():
# 		# If no such row exists, add the group name to the list
# 		missing_tlindex_groups.append(name)
# 
# # Output the results
# if missing_tlindex_groups:
# 	print("Groups missing TimeLapseIndex == 1:")
# 	for group_name in missing_tlindex_groups:
# 		print(group_name)
# else:
# 	print("All groups contain TimeLapseIndex == 1.")


## group the dataframe based on cell and treatment, 
## then calculate the percent change in area for each time point after treatment
area_result_df = uniqueID_df.groupby(['UniqueCellID', 'Treatment[mM]'], group_keys=False).apply(calculate_cellLevel_pcts, column=Area_columnName)

## trim down the columns to only those desired
area_result_df = area_result_df[columns_to_extract]
print(f'''the pct result df is:\n{area_result_df}''')

## group the dataframe based on cell and treatment, 
## then calculate the percent change in equivalent diameter for each time point after treatment
dia_result_df = uniqueID_df.groupby(['UniqueCellID', 'Treatment[mM]'], group_keys=False).apply(calculate_cellLevel_pcts, column=EqDia_columnName)

## trim down the columns to only those desired
dia_result_df = dia_result_df[columns_to_extract]
print(f'''the pct result df is:\n{dia_result_df}''')

## take only the first row so each cell for each treatment 
## is only represent once in the dataframe
a_result_df_first_row = area_result_df.groupby(['UniqueCellID', 'Treatment[mM]']).first().reset_index()
# print(f'''the first row for pct each group is:\n{a_result_df_first_row}''')
d_result_df_first_row = dia_result_df.groupby(['UniqueCellID', 'Treatment[mM]']).first().reset_index()
# print(f'''the first row for pct each group is:\n{d_result_df_first_row}''')

## pull out the replicate ID (aka GroupID) from the unique cell identifier 
## for easier reference and when making the replicate average dataframe
a_result_df_first_row['GroupID'] = a_result_df_first_row['UniqueCellID'].str.extract(r'([A-Z])')
# print(f'''the grouped rep pct df is:\n{a_result_df_first_row}\n\n''')
a_avg_pcts_df = get_rep_avgs(a_result_df_first_row, avging_groups_pcts, cellLevel_pcts)

d_result_df_first_row['GroupID'] = d_result_df_first_row['UniqueCellID'].str.extract(r'([A-Z])')
# print(f'''the grouped rep pct df is:\n{d_result_df_first_row}\n\n''')
d_avg_pcts_df = get_rep_avgs(d_result_df_first_row, avging_groups_pcts, cellLevel_pcts)

## for each measurement column, pivot the dataframe for easy import into prism
## each row is a cell (or replicate average) and each column is a treatment
## values will be based on the indicated measurement column
## saves pivoted dataframes to given output directory as .csv
for pct in cellLevel_pcts:
	## pivot and save area cell level data
	a_pivoted_df_pcts = pivot_df(a_result_df_first_row, 'UniqueCellID', 'Treatment[mM]', pct)
	save_to_csv(a_pivoted_df_pcts, output_directory, pct , 'cell_level_area')
	
	## pivot and save cell level data
	d_pivoted_df_pcts = pivot_df(d_result_df_first_row, 'UniqueCellID', 'Treatment[mM]', pct)
	save_to_csv(d_pivoted_df_pcts, output_directory, pct , 'cell_level_dia')

	## pivot and save area avgs
	a_pivoted_avgs_pcts = pivot_df(a_avg_pcts_df, 'GroupID', 'Treatment[mM]', pct)
	save_to_csv(a_pivoted_avgs_pcts, output_directory, pct , 'avg_area')
	
	## pivot and save dia avgs
	d_pivoted_avgs_pcts = pivot_df(d_avg_pcts_df, 'GroupID', 'Treatment[mM]', pct)
	save_to_csv(d_pivoted_avgs_pcts, output_directory, pct , 'avg_dia')
```
