## Supplemental files 1-3 for "Frog-killing chytrid fungi deploy different strategies to regulate intracellular pressure in cell types that have or lack a cell wall": 2025_Prostak_Osmoregulation_File_S3.py

```
## will take ga3 data for long term turgor sorbitol experiments
## adds replicate ID and treatment column based on file name
## will calculate the % change in EqDia across each frame in comparison to the before treatment per cell
## will format the data for easy input into Prism
## produces averages for each replicate too

## input document must be a .csv with the following columns:
	##FileName	TrackId	MultiPointIndex	TimeLapseIndex	Circularity	ObjectId	EqDiameter[um]
	## FileName must be in the following format: [string]_[YYYYMMDD]_[string]_###mM_###.nd2


import pandas as pd
from datetime import datetime


def get_uniqueCellID(row):
	'''
	Combines values for a cell to create a unique identification string for ease of reference and data handling
	Will extract the date, treatment, and file number from the file name for the given row
	
	Parameters:
		row (pd.DataFrame): a row of the filtered dataframe of interest
		
	Returns:
		 a string uniquely representing the cell in the given row 

	'''

	date = row['Date']
	treatment = str(row['Treatment(mM)'])
	file_ending = row['FileName'].split('_')[-1]
	file_num = file_ending.split('.')[0]
	rep = row['ReplicateID']
	cell_num = row['TrackId']
	return f'''{date}_{treatment}_{file_num}_{rep}_{cell_num}'''

def collapse_cellID_info(cellLevel_df, new_column_name, columns_to_remove, new_column_order):
	'''
	Adds a cell's unique identification string to a new column, removes unwanted columns, and reorders columns as desired.

	Parameters:
		cellLevel_df (pd.DataFrame): the dataframe of objects after filtering out non-analyzable objects
		new_column_name (str): the name of the column of ID strings
		columns_to_remove (list): a list of column names of columns to be removed
		new_column_order (list): a list of column names in the desired order
		
	Returns:
		cellLevel_df (pd.DataFrame): the input dataframe of objects, with the new column added, 
			unwanted columns removed, and columns in the desired order
	'''

	cellLevel_df[new_column_name] = cellLevel_df.apply(get_uniqueCellID, axis=1)
	cellLevel_df = cellLevel_df.drop(columns=columns_to_remove)
	cellLevel_df = cellLevel_df[new_column_order]
# 	print(f'''\n\nThe cell level data with unique cellIDs are:\n{cellLevel_df}''')
	return cellLevel_df

def save_to_csv(dataframe, output_directory, measure, level):
	'''
	Saves a dataframe as a .csv to a given directory, saving the file name using the inputed variables
	
	Parameters:
		dataframe (pd.DataFrame): the dataframe to be saved
		output_directory (str): the path to the directory in which the file is to be saved
			must NOT have ending /
		measure (str): the measure being saved (ex., "ratio2-1")
		level (str): the level of the measurement being saved (i.e., cell level, replicate average)
		
	Returns:
		None, writes a .csv to indicated directory
	'''
	
	file_path = f'''{output_directory}/{current_date}_{measure}_{level}.csv'''
	dataframe.to_csv(file_path, index=False)
	print(f'''The output for {measure}, {level} has been written to:\n{file_path}\n\n\n''')
	return None
	
	
def reorder_cell_level(df):
	'''
	
	Reorder the cell level dataframe to put the last two columns first
	Reordered dataframe will be used to pivot for desired result for import into prism

	Parameters:
		df (pd.DataFrame): the dataframe to be reordered
		
	Returns:
		df_reordered (pd.DataFrame): the dataframe where the last two columns 
			of the original dataframe are now the first two

	'''

	# Get the list of current columns
	columns = list(df.columns)

	# Extract the last two columns and the rest of the columns
	last_two_columns = columns[-2:]  # ['Treatment(mM)', 'ReplicateID']
	other_columns = columns[:-2]      # All other columns except the last two

	# Define the new column order by placing the last two columns at the front
	new_column_order = last_two_columns + other_columns

	# Reorder the DataFrame
	df_reordered = df[new_column_order].T
	return df_reordered
	

##########################################################################################
######################### BEGIN USER DEFINED INPUT #######################################

## No change needed: sets current date of when the script is run
current_date = datetime.now().strftime('%Y-%m-%d')

## Path to input .ga3 results file
ga3_file = "/Users/laboratory/Documents/Fritz_Laylin_Lab/Bd_osmoreg/turgor/Sorb/GA3_output/LT-sorb_GA3_v3_output_run_11-18-24.csv"

## Path to DIRECTORY to save output files to-- 
## individual files will be saved with unique file names according to 
## current date, measurement type, and cell vs replicate level
## DO NOT include the final "/" in the directory
output_directory = '/Users/laboratory/Documents/Fritz_Laylin_Lab/Bd_osmoreg/turgor/Sorb/replicates_for_prism/long-term_sorb/replicates'

## the number of frames an object needs to be present for analysis
total_frames = 25

## the area on the lower limit of the distribution to remove objects that are likely improperly recognized cells
## in um^2, as an integer
lower_area_cutoff = 130

## the area on the upper limit of the distribution to remove objects that are likely multiple cells
## in um^2, as an integer
upper_area_cutoff = 500


## Dictionary mapping the dates from the files to a replicate ID number for ease of reference
rep_dict = {"20240925":"A", "20240926":"B", "20240927":"C"} 

## List of the order of columns for the table after when making unique cell IDs
new_column_order = ['UniqueCellID', 'TimeLapseIndex', 'ReplicateID', 'Treatment(mM)', 'BinEqDiameter[um]', 'BinObjectArea[um2]', 'time(min.)']

new_column_name = "UniqueCellID" 

## List of columns to remove after creating a unique cellID column
columns_to_remove = ['FileName', 'BinCircularity', 'ObjectId']


######################### END USER DEFINED INPUT #########################################
##########################################################################################

## import the ga3 data into a pandas dataframe
og_df = pd.read_csv(ga3_file)
# print(og_df)


## pull out the date associated with the file, put it in a new column
og_df['Date'] = og_df['FileName'].str.extract(r'_(\d{8})_')

## assign each date a replicate ID for ease of reference, add new column with these values
og_df['ReplicateID'] = og_df['Date'].map(rep_dict)

## pull the treatment amount from the file name, add it to a new column 
og_df['Treatment(mM)'] = og_df['FileName'].str.extract(r'_(\d+)mM_')

# print(og_df)


## group the data by file name and trackID so all frames for one cell stay together 
grouped_df = og_df.groupby(['FileName', 'TrackId'])
# print(grouped_df)


## Filter out cells that do not fit frame number or area threshold for analysis
filtered_df = grouped_df.filter(
    lambda group: (
        group['TimeLapseIndex'].count() == total_frames and  # Ensure all frames are present
        all((lower_area_cutoff <= group['BinObjectArea[um2]']) & (group['BinObjectArea[um2]'] <= upper_area_cutoff)) # Exclude groups where at least one value is outside of the cutoff range
    )
)


## add a new column converting the time lapse index (frame number) into minutes
## frame 1 is 0 min, frame 2 is 5 min, and so on based on imaging parameters
filtered_df['time(min.)'] = (filtered_df['TimeLapseIndex'] - 1) * 5
# print(filtered_df)

## Check the number of cells that made it through filtering
total_rows = filtered_df.shape[0]
total_cells = total_rows/total_frames
print(f'''\n\n\nThe total number of cells analyzed is: {total_cells}\n\n\n''')

## create and add the unique identification strings for the cells, remove unwanted columns, reorder columns
uniqueID_df = collapse_cellID_info(filtered_df, new_column_name, columns_to_remove, new_column_order)
print(uniqueID_df)

## save the dataframe of filtered cells with their uniqueIDs
save_to_csv(uniqueID_df, output_directory, 'filtered-data', 'UniqueIDs')


# Create an empty dictionary to store the normalized results
normalized_eqDia = {}
normalized_area = {}


# Group by UniqueCellID
grouped = uniqueID_df.groupby('UniqueCellID')

# Iterate over each group (cell)
for name, group in grouped:
	
	# Take the first value in BinEqDiameter[um] as the initial diameter
	d_initial = group['BinEqDiameter[um]'].iloc[0]
    
	# Normalize the BinEqDiameter[um] values by dividing by the initial diameter, turn it into a percent
	d_normalized = (group['BinEqDiameter[um]'] / d_initial) * 100
    
	# Use time(min.) as the key for this group, add group to normalized dictionary
	normalized_eqDia[name] = pd.Series(d_normalized.values, index=group['time(min.)'].values)
	
	
	# Take the first value in BinObjectArea[um2] as the initial area
	a_initial = group['BinObjectArea[um2]'].iloc[0]
	
	# Normalize the BinEqDiameter[um] values by dividing by the initial area, turn it into a percent
	a_normalized = (group['BinObjectArea[um2]'] / a_initial) * 100

	# Use time(min.) as the key for this group, add group to normalized dictionary
	normalized_area[name] = pd.Series(a_normalized.values, index=group['time(min.)'].values)


# Convert the dictionary to a DataFrame
d_normalized_df = pd.DataFrame(normalized_eqDia)
a_normalized_df = pd.DataFrame(normalized_area)


# Transpose the DataFrame to make each row a UniqueCellID and columns as calculated values
d_normalized_df = d_normalized_df.T
a_normalized_df = a_normalized_df.T

# print(d_normalized_df)
# print(a_normalized_df)


# Extract ReplicateID and Treatment(mM) for each UniqueCellID from the original DataFrame
d_normalized_df['ReplicateID'] = uniqueID_df.groupby('UniqueCellID')['ReplicateID'].first().values
d_normalized_df['Treatment(mM)'] = uniqueID_df.groupby('UniqueCellID')['Treatment(mM)'].first().values

a_normalized_df['ReplicateID'] = uniqueID_df.groupby('UniqueCellID')['ReplicateID'].first().values
a_normalized_df['Treatment(mM)'] = uniqueID_df.groupby('UniqueCellID')['Treatment(mM)'].first().values

## Reorder the columns so ReplicateID and Treatment(mM) come first
d_normalized_df_reordered = reorder_cell_level(d_normalized_df) 

a_normalized_df_reordered = reorder_cell_level(a_normalized_df)
# print(a_normalized_df_reordered)


# Reset the index to turn 'Treatment(mM)' and 'ReplicateID' from index to row names
save_to_csv(d_normalized_df_reordered.reset_index(), output_directory, "PctsEqDia", "cellLevel")
save_to_csv(a_normalized_df_reordered.reset_index(), output_directory, "PctsArea", "cellLevel")


# Group by Treatment(mM) and ReplicateID, then calculate the mean and stdev for each time point
d_grouped_mean = d_normalized_df.groupby(['Treatment(mM)', 'ReplicateID']).mean()
d_grouped_stdv = d_normalized_df.groupby(['Treatment(mM)', 'ReplicateID']).std()

a_grouped_mean = a_normalized_df.groupby(['Treatment(mM)', 'ReplicateID']).mean()
a_grouped_stdv = a_normalized_df.groupby(['Treatment(mM)', 'ReplicateID']).std()

# print(f'''BEFORE:\n\n{grouped_mean}\n\n''')
# print(grouped_stdv)

# Reset the index to turn 'Treatment(mM)' and 'ReplicateID' from index to row names
d_grouped_mean_T = d_grouped_mean.T
d_grouped_stdv_T = d_grouped_stdv.T

a_grouped_mean_T = a_grouped_mean.T
a_grouped_stdv_T = a_grouped_stdv.T

## save the dataframes
save_to_csv(d_grouped_mean_T.reset_index(), output_directory, "PctsEqDia", "Avgs")
save_to_csv(d_grouped_stdv_T.reset_index(), output_directory, "PctsEqDia", "Stdvs")

save_to_csv(a_grouped_mean_T.reset_index(), output_directory, "PctsArea", "Avgs")
save_to_csv(a_grouped_stdv_T.reset_index(), output_directory, "PctsArea", "Stdvs")

print(f'''\n~~~~~~~~~~DONE~~~~~~~~~~\n''')
```
